# Interpreting the Effects of DNA Polymerase Variants at the Structural Level Using MAVISp and Molecular Dynamics Simulations

**DOI:** 10.1101/2025.08.04.653638

**Authors:** Matteo Arnaudi, Karolina Krzesińska, Ludovica Beltrame, Pablo Sánchez-Izquierdo Besora, Matteo Tiberti, Mef Nilbert, Anna Rohlin, Elena Papaleo

## Abstract

Genetic variants in the DNA polymerase enzymes POLE and POLD1 can affect protein function by altering stability, catalysis, DNA binding, and interactions with other biomolecules. Understanding the structural basis of these variants is important for a comprehensive interpretation of variant impact. In this study, we used MAVISp, a modular structure-based framework, and molecular dynamics simulations to analyze over 60,000 missense variants of POLE and POLD1. By integrating results from changes in folding and binding free energies, local alterations in the proximity of the active or phosphorylation sites, we provided a detailed structural interpretation of variants reported across various databases, including ClinVar, COSMIC, and cBioPortal. Moreover, we predicted the functional consequences of variants not found yet in disease-related databases, thereby creating a comprehensive catalogue for future studies. Of note, our approach enabled us to classify 364 Variants of Uncertain Significance (VUS) as PP3 evidence and 323 as BP4 evidence, in accordance with the American College of Medical Genetics and Genomics (ACMG) guidelines. Additionally, we identified a group of variants that could alter the native orientation of the residues within the catalytic site of the exonuclease domain, such as POLE variants P297S and P436R. Finally, we identified a group of variants predicted to affect DNA-binding affinity and rationalized their effects in terms of different energetic contributions and structural features. Collectively, our results not only advance our understanding of protein variant effects in POLE and POLD1 at the structural level but also support future studies aimed at variant classification, variant prioritization for experimental studies, and functional interpretation across diverse biological contexts.

## Introduction

DNA polymerases ε (POLE) and δ (POLD1) play important roles in DNA replication and ensure replication fidelity through their base selectivity and proofreading activities [1]. Genetic variants in POLE and POLD1 could affect their structural integrity and function, potentially altering the accuracy of DNA replication [2]. For example, variants affecting POLE and POLD1 have been implicated in conditions characterized by increased genomic instability [3].

POLE is involved in the replication of the leading strand of DNA, while POLD1 synthesizes the Okazaki fragments of the lagging strand of genomic DNA [4,5]. The exonuclease domains (residues 268-472 in POLE, and residues 309-526 in POLD1) are primarily responsible for the correction of rare errors that might occur during the DNA replication process [1]. The proofreading activity ensures high-fidelity replication, with remarkably low error rates of 10^-7^-10^-8^ errors per base pair [6]. Moreover, the two enzymes interact with several accessory subunits, which ensure their stability and functionality, and operate in concert with other proteins to form the complex replisome machinery.

Our group recently proposed a structure-based framework, MAVISp (Multi-layered Assessment of Variants by Structure for proteins), to characterize and classify missense variants found in ClinVar [7] and other cancer datasets in a high-throughput manner [8]. MAVISp is based on the principle that mutations in coding regions can have diverse consequences for the resulting protein and its cellular function. These may include changes in structural stability, altered interactions with other proteins or biomolecules (e.g., DNA), disruption of catalytic or cofactor-binding sites, and alterations in post-translational modifications. The MAVISp framework also annotates variants from disease databases, predicts their effect using different Variant Effect Predictors (VEPs), and finally uses structure-based methods to identify potential structural mechanisms of missense mutations, namely, mechanistic indicators. The MAVISp protocol can be applied in both the so-called *simple* and *ensemble modes*. The *ensemble mode* accounts for protein dynamics and, for instance, leverages structures from molecular dynamics (MD) simulations.

In this study, we collected new data not previously available in the MAVISp database, applying most of the modules implemented in the MAVISp toolkit. We also performed additional analyses of MD simulations, to systematically characterize all possible missense mutations in the folded regions of POLE and POLD1. Additionally, we interpreted the results on a case-by-case basis in a contextualized manner. In fact, MAVISp can be applied to very diverse proteins, and the results from VEPs should be interpreted in the context of the specific biological and structural features of the protein, and in relation to the available information on the phenotype associated with the variant. This consideration is especially important for multidomain proteins, where mutations in distinct domains can give rise to distinct phenotypes. The DNA polymerases POLE and POLD1 are good examples for which the functional impact of mutations is highly dependent on the biological context.

The role of POLE and POLD1 as tumor suppressors remains poorly defined. Mutations in the exonuclease (proofreading) domain, known to impair replication fidelity and to promote a tumorigenic phenotype [9], are classified as loss-of-proofreading-function in hereditary cancer syndromes and hypermutated tumors. For this type of variant, a damaging prediction from VEPs is likely to reflect a relevant biological effect and may be useful for variant prioritization for further studies. In contrast, damaging predictions from VEPs for variants located in the polymerase domain should be interpreted with caution since an intact activity of the polymerase domain is important to support malignant proliferation [10,11]. Since polymerase activity is required for cell viability in cancer cells, these variants are unlikely to cause a complete loss of protein function but are more likely to impair replication fidelity. In these cases, a detailed structural analysis is necessary to properly assess the functional impact of the variant and its relevance to the specific phenotype under investigation. In general, given that such predictions are context-dependent, variants that appear poorly informative in clinical settings could still have relevance in other contexts. For example, amino acid substitutions in the polymerase domain that are predicted to impair function may provide useful insights for protein engineering applications aimed at modulating polymerase activity.

Overall, by integrating saturation mutagenesis data with structural, functional, and interaction-based analyses, we aim to identify possible molecular mechanisms underlying both known (likely) pathogenic and previously uncharacterized variants in POLE and POLD1, thereby building a structural-level atlas of variant effects. In addition, we developed new protocols for analyses in the MAVISp *ensemble mode* to elucidate the effects of variants predicted to destabilize the native architecture of active or cofactor binding sites in a conformation-dependent manner, as well as to analyze variant sites with low confidence in the initial structure that may reflect underlying structural flexibility when protein dynamics is taken into account. Finally, we proposed a protocol within MAVISp to classify VUS in the exonuclease domain of both proteins, according to the American College of Medical Genetics and Genomics (ACMG) criteria, leveraging the protein-specific data aggregated through the MAVISp workflow, drawing inspiration from a recently published study [12].

## Results and Discussion

### Overview of variants characterized in the study and structure selection

To orient the reader, we first summarize the main sources of variants included in the study and the protein structures and complexes used for analysis. Using an *in-silico* saturation mutagenesis approach, we generated data for 43415 variants in POLE (residues 2-2286, **Figure S1A**) and 21014 variants in POLD1 (residues 2-1107, **Figure S1B**). Among these, 4291 POLE (**Figure S1C**) and 2025 POLD1 (**Figure S1D**) variants were reported in ClinVar. Of those, 1966 variants in POLE and 1990 variants in POLD1 have been annotated in ClinVar as associated with colorectal cancer, polymerase proofreading-associated polyposis (PPAP), or hereditary cancer-predisposing syndrome (**see OSF repository**, *variants_output.csv*). The three-dimensional (3D) models shown in **Figure S1A** and **S1B** for POLE and POLD1, respectively, were used as input structures to analyze the effects of variants on structural stability, functional sites, long-range interactions, and phosphorylation.

As previously mentioned, POLE and POLD1 are the catalytic subunits of the DNA replication machinery, responsible for leading- and lagging-strand elongation, respectively. Their polymerase activity is regulated through interactions with several subunits of the replication machinery, which are essential for enzymatic processivity. Additionally, the replication machinery can contribute to multiple DNA repair pathways [13] [14] [15]. For these reasons, we analyzed the effects of variants on the protein-protein complexes involving POLE and five interactors (PCNA, CDC45, POLE2, MCM2, and MCM5), as well as POLD1 and four interactors (PCNA, POLD4, POLD2, and LASP1). As detailed in the Methods, these interactors were selected because they were the only ones for which the 3D structures of their protein–protein complexes met the quality-control criteria of the MAVISp protocol. Furthermore, we assessed the impact of amino acid substitutions on DNA interaction for both POLE and POLD1. To this end, we used experimentally determined 3D structures of POLE and POLD1 in complex with DNA, as well as five additional POLE 3D structures representing key intermediates in base-pair mismatch recognition and excision (**Materials and Methods**). We applied both the *simple* and *ensemble modes* of MAVISp. The only exceptions were the analyses of variants in the proximity of functional sites, which are currently implemented only in the *simple mode*, and analyses of variant effects on local interactions, which were not performed in the *ensemble mode* due to computational time constraints. Unless otherwise specified, the results discussed in the text refer to the *ensemble mode*, as it captures the dynamic nature of protein structures. Where available, we also report experimental data for variants (as detailed in the **Materials and Methods Section**).

In **Figure 1**, we illustrate a schematic workflow for the calculations and analyses performed in this study, distinguishing between the application of MAVISp to known pathogenic variants, where we only search for mechanistic indicators at the structural level, and applications to characterized VUS or other variants not yet reported in ClinVar.

**Figure 1.**
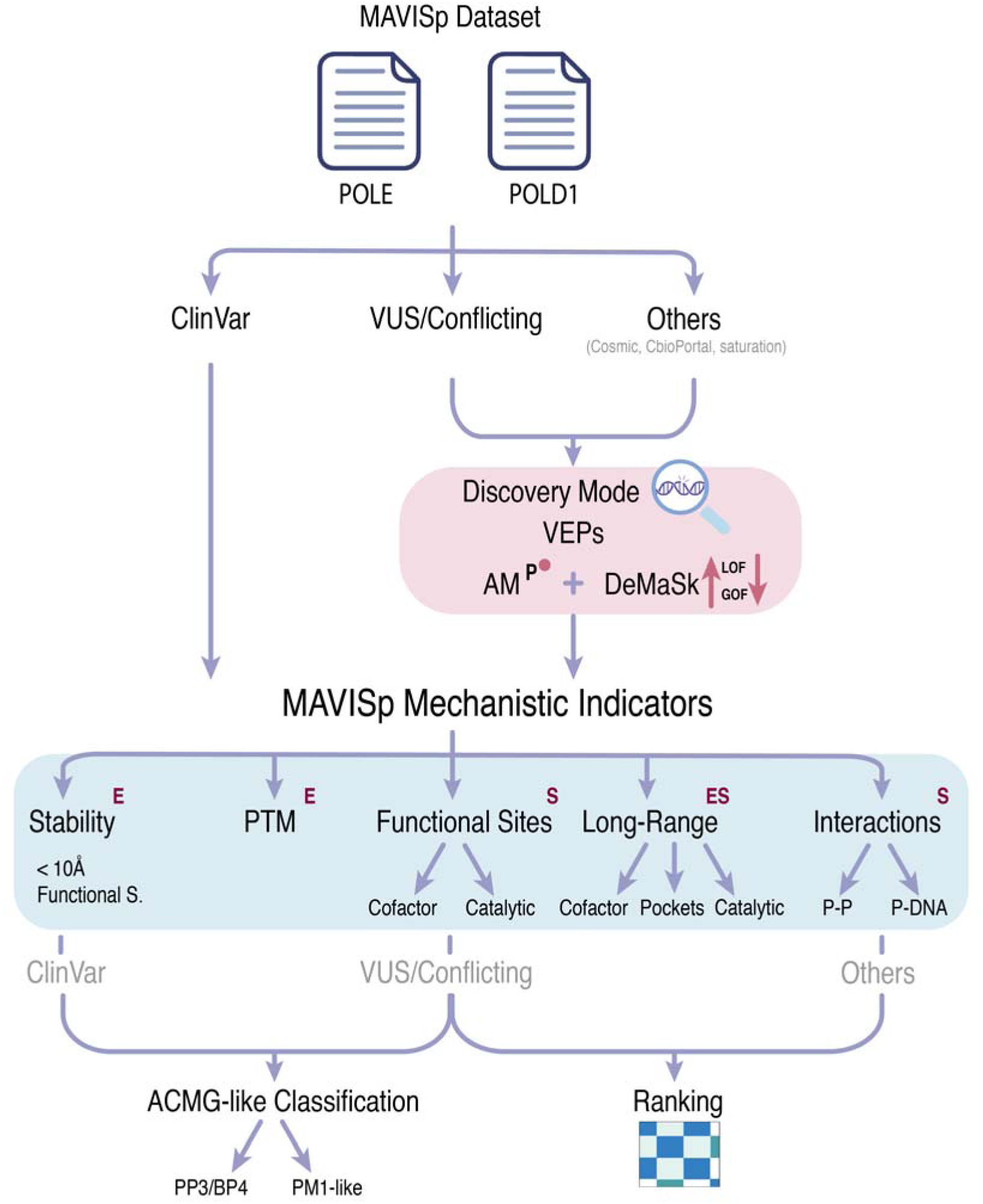
Analysis pipeline applied in this study using VEPs and MAVISp mechanistic indicators. The figure illustrates the analysis pipeline that we generally employ for focused studies, which we have further expanded in this work to include variant ranking and ACMG-like classifications. Following protein curation and data collection with MAVISp, the variants within the selected protein structure are categorized, according to their mutation source, into known ClinVar pathogenic variants, VUS/Conflicting, and Others, which may include variant sources from CBioPortal, COSMIC, and saturation. The VUS/Conflicting and Others groups undergo a variant discovery mode, where we filter for variants with an AlphaMissense predicted ‘pathogenic’ annotation and with a DeMaSk loss or gain of fitness (LOF and GOF, respectively) annotation. Subsequently, across all variant groups, we investigated the identified MAVISp mechanistic indicators in *simple mode* (S), *ensemble mode* (E), or both. The data collected with MAVISp for the known ClinVar pathogenic and benign variants can also be used to calibrate a classifier to apply to ClinVar VUS/Conflicting variants according to the American College of Medical Genetics and Genomics (ACMG) guidelines in PP3/BP4 classes and providing PM1-like evidence. While, for VUS/Conflicting and Other groups of variants, a feature-based ranking pipeline is also applied using MAVISp structural evidence to identify and prioritize variants.

### Performance evaluation of MAVISp VEPs used to identify potentially pathogenic variants

Before engaging in the search for mechanistic indicators, we assessed how effectively the VEPs integrated into MAVISp and used as the first layer of assessment in the discovery workflow can identify disease-relevant variants in POLE and POLD1. As illustrated in **Figure 1**, the initial step of the default MAVISp discovery workflow for evaluating variants of uncertain significance (VUS) or other unclassified variants not yet reported in ClinVar is to prioritize variants predicted as damaging by the VEP AlphaMissense [16]. This choice is motivated, on one hand, by the fact that the MAVISp framework is rooted in structural methods, and, on the other hand, by the extensive coverage currently provided by AlphaMissense for proteins included in the MAVISp database. Nevertheless, when focusing on a specific group of proteins, it is important to evaluate whether modifications to the discovery workflow could offer advantages. In this context, additional VEPs are integrated into the MAVISp framework and can guide the first step of the discovery workflow, such as EVE [17], REVEL [18], or GEMME [19].

We identified 37 (POLE) and 12 (POLD1) ClinVar-classified variants in the MAVISp datasets with available clinical annotations and coverage across all the VEPs (benign, likely benign, pathogenic, or likely pathogenic; see **Table S1)**. The number of well-characterized variants for each gene is limited; therefore, we pooled data from both proteins for the analysis reported in **Table 1**. Because the datasets are unbalanced, with a predominance of benign variants, we focused primarily on the Matthews Correlation Coefficient (MCC) and balanced accuracy. As shown in **Table 1**, AlphaMissense achieved the best-balanced accuracy, whereas EVE yielded the highest MCC score [16,17]. In addition, by examining the complete MAVISp datasets for POLE and POLD1, we observed that AlphaMissense provides full coverage (100%) across POLE and POLD1 variants compared to EVE (78.68 %), highlighting its general utility in the MAVISp discovery workflow for this specific study.

**Table 1.**
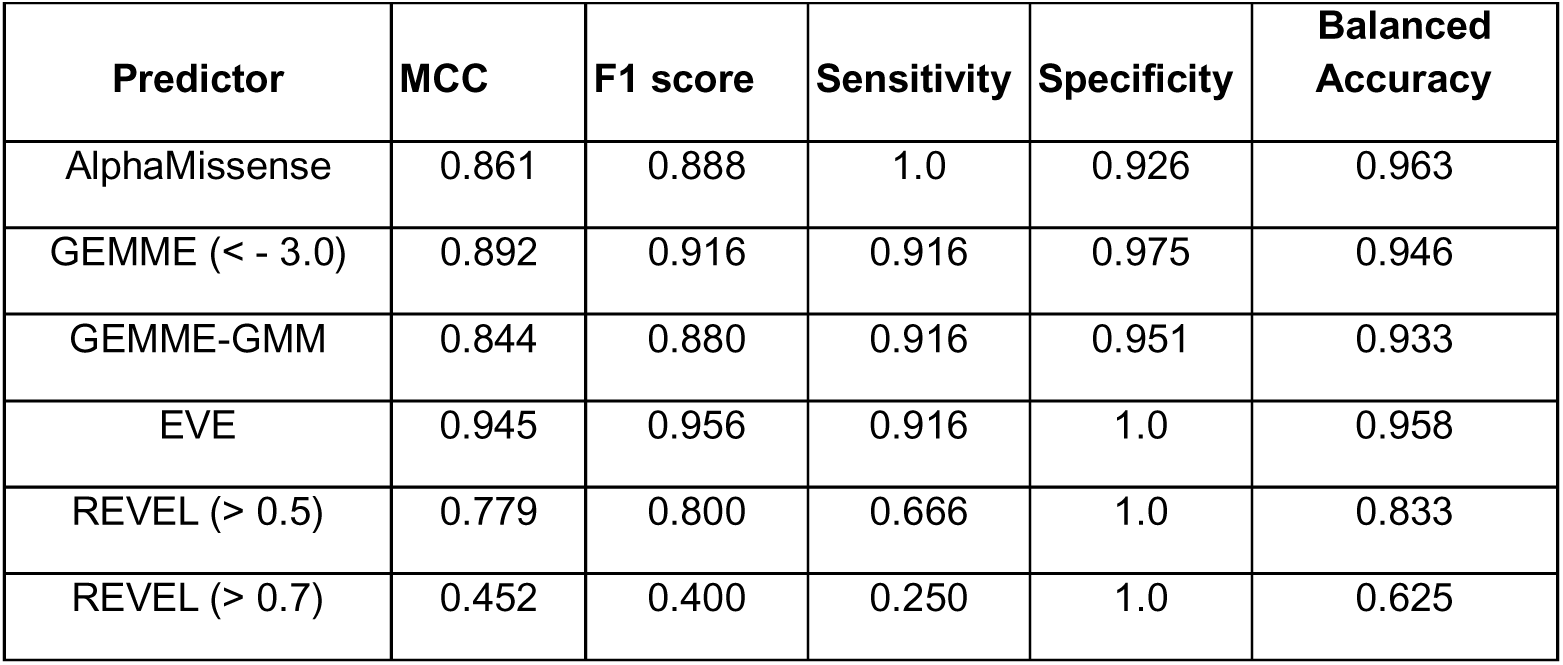
Performance metrics of the VEPs available in MAVISp for POLE and POLD1. Here we have only considered variants that were reported in ClinVar as pathogenic or benign, and used them as ground truth to evaluate the performance of the VEPs (EVE, AlphaMissense, GEMME and REVEL). It should be noted that MAVISp also supports DeMaSk as VEP, but it has not been included in this comparison. This is because we use DeMaSk as a second layer of evidence to take advantage of its gain/loss-of-fitness classification and because it has been trained on experimental deep mutational data, as detailed in **Figure 2**.

| Predictor | MCC | F1 score | Sensitivity | Specificity | Balanced Accuracy |
| --- | --- | --- | --- | --- | --- |
| AlphaMissense | 0.861 | 0.888 | 1.0 | 0.926 | 0.963 |
| GEMME (< - 3.0) | 0.892 | 0.916 | 0.916 | 0.975 | 0.946 |
| GEMME-GMM | 0.844 | 0.880 | 0.916 | 0.951 | 0.933 |
| EVE | 0.945 | 0.956 | 0.916 | 1.0 | 0.958 |
| REVEL (> 0.5) | 0.779 | 0.800 | 0.666 | 1.0 | 0.833 |
| REVEL (> 0.7) | 0.452 | 0.400 | 0.250 | 1.0 | 0.625 |

### Structural mechanisms in pathogenic and likely pathogenic variants of POLE and POLD1

One application of the MAVISp toolkit is to identify the structural mechanisms underlying variants in proteins of interest that are already reported as pathogenic or likely pathogenic in ClinVar. These analyses are useful to acquire new insights on the effects exerted by these variants that can guide, for example, experimental validation or inform on their mechanism of action. We therefore retained only variants classified as pathogenic or likely pathogenic in ClinVar and for which at least one mechanistic indicator was provided by MAVISp (**Figure 1**). To achieve this goal, we leveraged the downstream analysis toolkit of MAVISp (**Materials and Methods**) using dotplots for a first data exploration to facilitate the identification of variants for a more in-depth investigation (**Figure 2A-B)**. Overall, MAVISp enabled us to provide structural insights for the variants S297F (p.Ser297Phe), N336K (p.Asn363Lys), P436R (p.Pro436Arg), of POLE and L474P (p.Leu474Pro) of POLD1 (see next Result sections).

**Figure 2.**
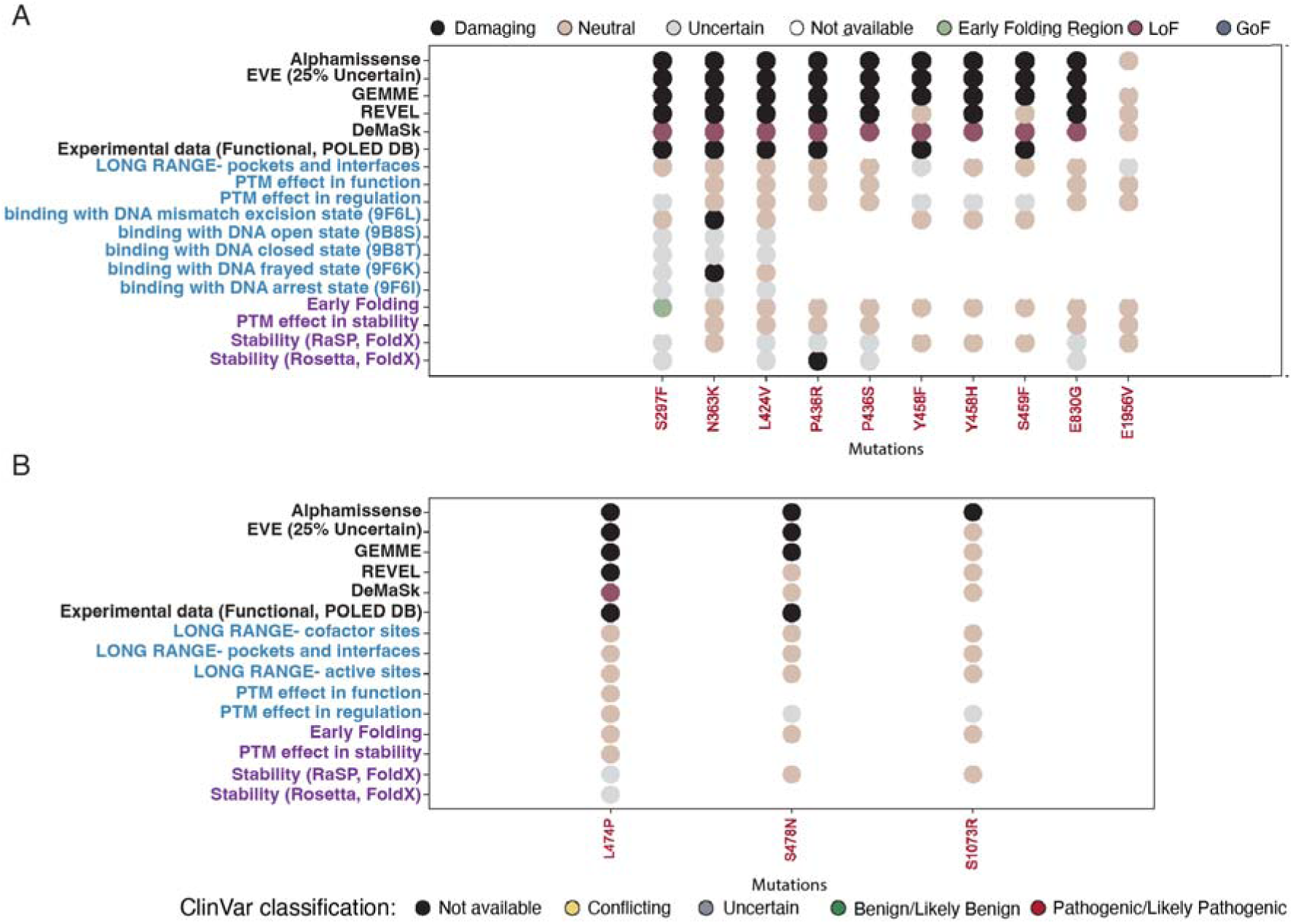
Analyses of structural mechanistic indicators for known pathogenic and likely pathogenic variants of POLE and POLD1 with MAVISp. We used dot-plot representation to get a first overview of the results from the different MAVISp modules in a simplified manner for POLE (A) and POLD1 (B), focusing only on the variants labelled as pathogenic or likely pathogenic in ClinVar (red colors on the x-axis). In the MAVISp dotplot each column corresponds to a single variant, and each dot indicates a predicted effect or annotation derived from the MAVISp modules. MAVISp features associated with stability are shown in purple, whereas features related to different functional properties are shown in blue. Rows in light green report results from VEPs or available experimental data. We only report the results from mechanistic indicators or other modules that are relevant to the results for sake of clarity. The full version of the dotplots is available in the **OSF repository** associated with the publication (https://osf.io/z8x4j). The results from the LONG_RANGE classification, Stability Classification, and PTM effects refer to calculations performed in the *ensemble mode* of MAVISp, the rest in the *simple mode*. We can observe that N363K in POLE has conformation-dependent damaging effects on the structure in complex with DNA, and P346R in POLE has effects on structural stability. The stability classification is based on a consensus approach between FoldX and RaSP, using 25 structures extracted from the MD simulations and evaluating average changes in folding free energy upon amino acid substitution (see Materials and Methods). Additionally, several variants in POLE and L474P in POLD1 remain uncertain for structural stability, which is an aspect that can be further investigated to identify conformation-dependent effects (see panel B in Figure 3).

All the variants of interest for this part of the study were located in regions with a pLDDT score >= 70%, except for E1956V in POLE (**Figure 3A**, pLDDT = 29.84%). We thus examined how the mutation site 1956 behaves during POLE MD simulation in terms of structural flexibility. Indeed, a site with a low pLDDT score in the initial static model may correspond to a flexible region that exhibits conformational heterogeneity when protein dynamics is taken into account (**see Materials and Methods**). The site is classified as flexible (see Root Mean Square Fluctuation, RMSF, results in *OSF repository*) and lies within a loop of the C-terminal region involved in inter-subunit interactions in the early stages of DNA replication and essential for the enzyme processivity [20] [21] [22].

**Figure 3.**
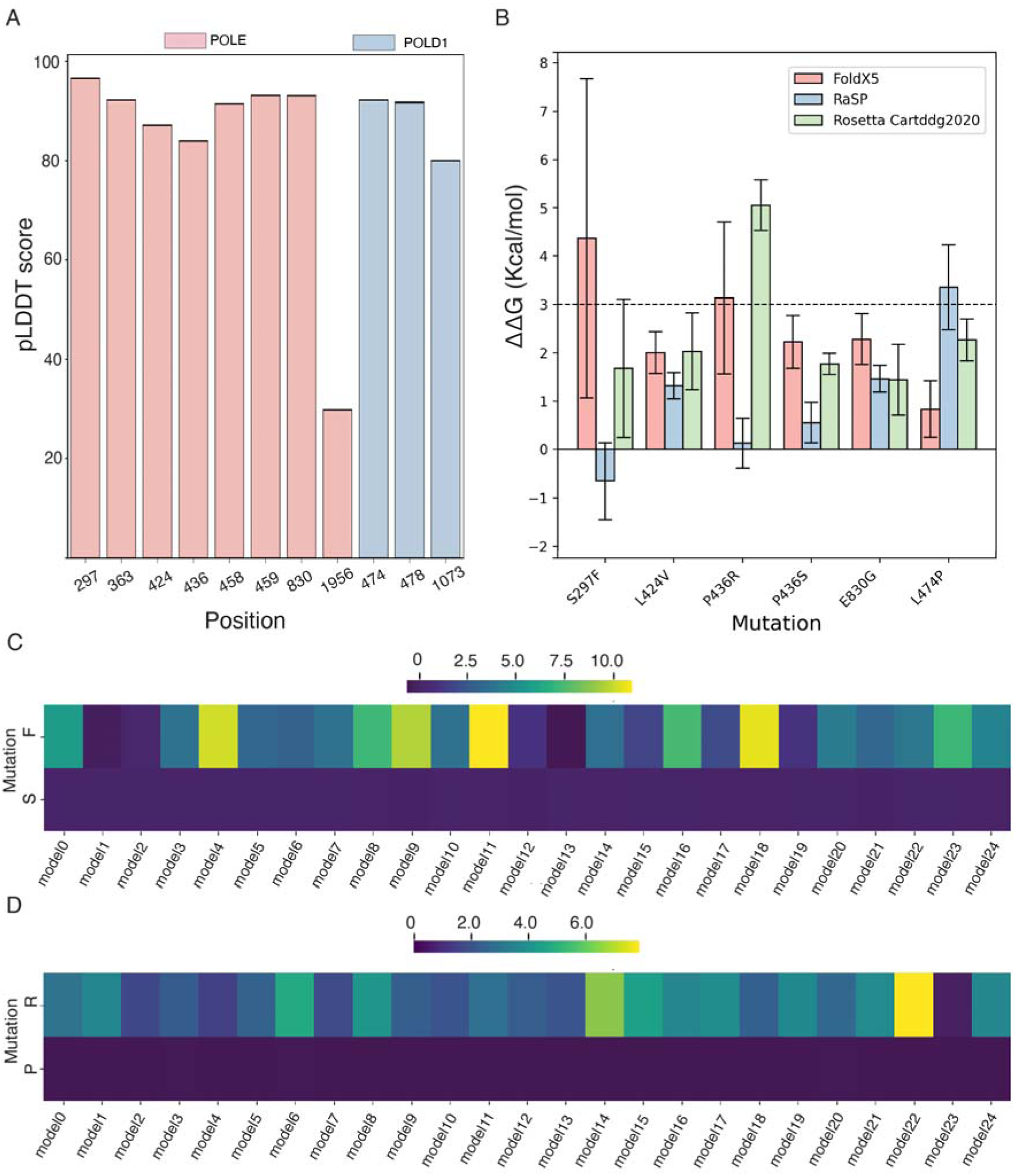
Analysis of the effects on structural stability in the proximity of the catalytic site of the exonuclease domain of POLE and POLD1 of pathogenic and likely pathogenic variants. (A) The bar plot shows the per-residue pLDDT scores for POLE (orange) and POLD1 (blue) at positions harboring ClinVar-reported pathogenic variants, providing an estimate of the local structural quality of the model. We can observe that E1956 in POLD1 is the only mutation site predicted with a pLDDT value lower than 70, which might also underlie an intrinsic flexibility of this region. We thus further verify the protein flexibility at this site during simulations using root-mean-square fluctuations as a flexibility probe (see Materials and Methods). (B) The bar plots report the average ΔΔG values predicted by RaSP, FoldX, and Rosetta, computed across 25 structures extracted from the MD simulations (RaSP, FoldX) or the three cluster centroids from the simulation upon structural clustering on the main-chain root-mean square deviation matrix (Rosetta). More details on these calculations are available in the Materials and Methods. Of note, RaSP prediction for S297F and P436R disagree with the rest of the predictors, which prompted us to a closer investigation on the structural effects exerted by these variants, collecting MD simulations of the variants themselves (see Figure 3). (C-D) The panels show the heatmaps in relation to FoldX-predicted changes of folding free energies across the 25 structures extracted from the MD simulations of POLE with focus on the position S297 (C) and P436 (D), and in relation to the changes induced by the S297F and P436R variants compared to the self-mutation assessment at the same position, which provides a control. Indeed, to assess the local quality of the structural models around positions 297 and 436 of POLE, we used a self-mutation analysis with FoldX (see Materials and Methods). This type of control is expected to result in minimal changes in folding free energy, typically close to 0 kcal/mol [23]. In our case, the predicted absolute ΔΔG values for the self-mutation at positions 297 and 436 were never below ∼0.5 kcal/mol, suggesting that the model used in this calculation is at an energy minimum.

Furthermore, we identified six known (likely) pathogenic variants with predicted uncertain effect on structural stability (**Figure 2A-B**) using the consensus method between FoldX and RaSP, namely S297F (p.Ser297Phe), L424V (p.Leu424Val), P436R/S (p.Pro346Arg/Ser), E830G (p.Glu830Gly) in POLE, and L474P (p.Leu474Pro) in POLD1. These uncertain predictions result from disagreement between the two methods used in the consensus approach (**Figure 1, and Materials and Methods**), as can be observed upon closer inspection of the predicted ΔΔG values (**Figure 3B**). The consensus between FoldX and RaSP constitutes the default protocol for assessing structural stability in MAVISp. Nevertheless, the framework also supports folding free energy calculations based on Rosetta. We therefore collected additional data using the Rosetta Cartesian protocol (**see Materials and Methods**) which provides more accurate predictions than its machine-learning-based alternative RaSP.

We could identify three classes of cases. In one hand, the POLD1 L474P variant showed marginal changes in ΔΔG according to FoldX but was predicted to be overall destabilizing by RaSP (**Figure 3B**). To further investigate this variant, we performed MD simulations of the mutant variant also considering that L474 is at the beginning of a short α-helix in proximity of the catalytic residue D515. We did not observe major changes in the protein conformation or in secondary structures that could support a destabilizing effect of this variant. Additionally, we observed only modest changes in contact occurrence, primarly attributable to differences in side-chain geometry and size between the leucine and proline residues (see **OSF repository**, contact analysis with CONAN for POLE and POLD1).

The second class of variants with uncertain effects on structural stability includes the POLE variants E830G, P436S, and L424V. For these variants, only one or two conformations among those used for FoldX calculations (i.e., < 10 %, see **OSF repository**, stability heatmaps) showed predicted destabilizing effects, suggesting that these variants are more likely to be neutral with respect to structural stability. This interpretation is further supported by both Rosetta and RaSP predictions (**Figure 3B**).

In contrast, in the third class of variants, i.e., the POLE variants S297F and P436R, we identified clear conformation-dependent effects, indicating a context-specific destabilization **(Figure 3C-D**). Specifically, 68% and 44% of the conformations used for FoldX calculations exhibited destabilizing effects, for S297F and P436R, respectively.

### S297F and P436R are known likely pathogenic POLE variants, which cause structural destabilization in the proximity of the catalytic site in the exonuclease domain

S297F is located in the exonuclease domain (**Figure 4A**) and is known to be associated with autosomal dominant POLE-related disorders [9], in which the cancer-prone phenotype is expected to require a functionally active POLE with reduced replication fidelity. This observation prompted us to speculate that the variant could cause localized destabilization at this site. Such partial destabilization may preserve the overall domain architecture while nonetheless impairing the enzymatic activity.

**Figure 4.**
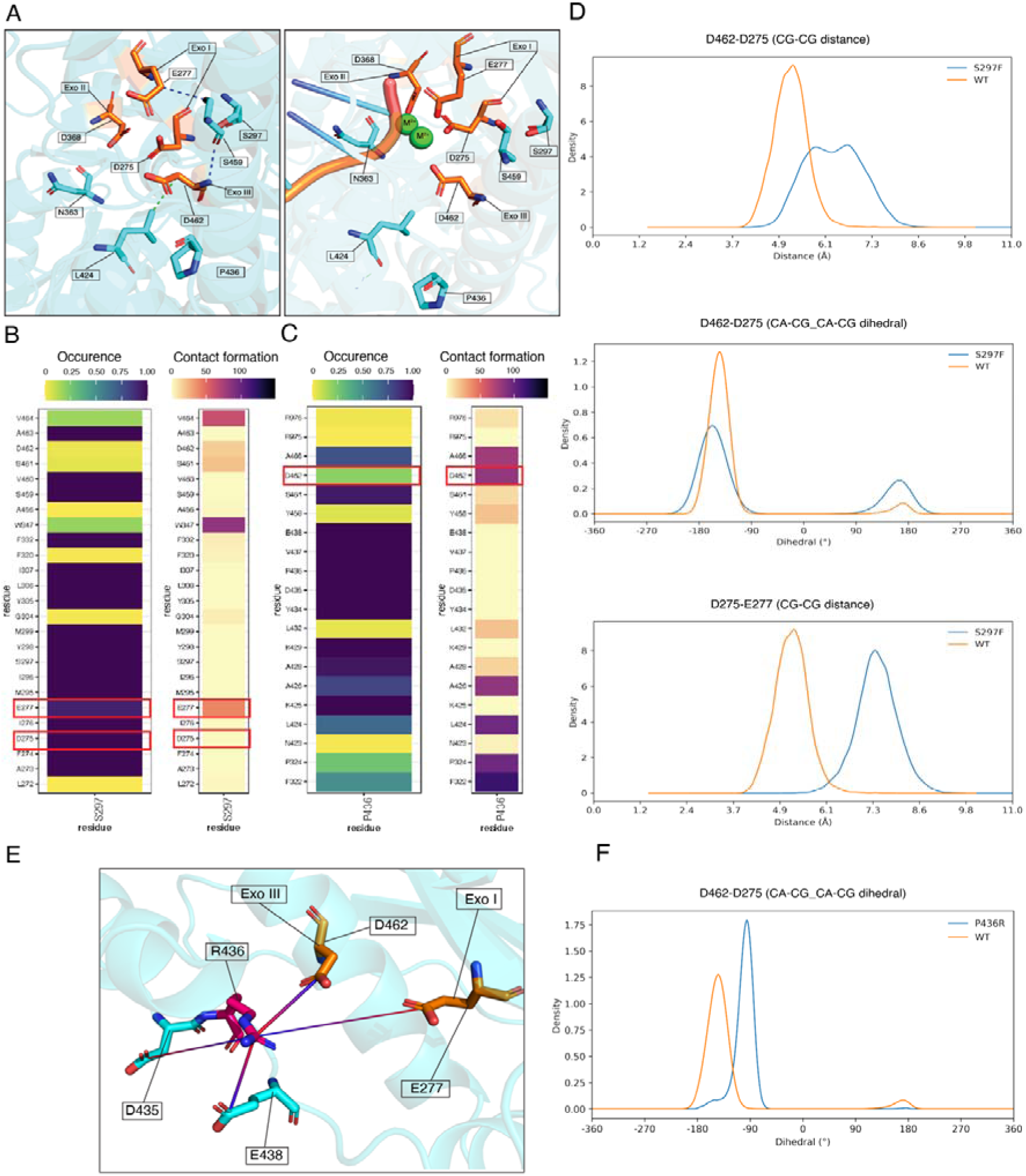
Structural investigation of the Effect of mutations reported Pathogenic in ClinVar on the exonuclease catalytic sites. A) Structural representation of the POLE exonuclease-domain catalytic center in the absence (left) and presence (right) of DNA in the mismatch-excision state (PDB ID: 9F6L). Catalytic residues are highlighted in orange, second-sphere residues in light blue. Key interactions are shown as hydrogen bonds (blue) and van der Waals contacts (green). B–C) Contact analysis of residues S297 and P436 along the POLE MD trajectory, indicating the occurrence and persistence of residue–residue contacts over time (contact frequency expressed as the number of frames in which the interaction is formed). D) Collective variables (CVs) extracted from MD simulations of the wild-type and S297F mutant, illustrating mutation-induced changes in distances and orientations of catalytic residues. Upper left: distance D275–E277; upper right: distance E277–D462; lower left: dihedral angle D275–D462; lower right: dihedral angle D275–E277. E) Structural representation of the salt bridges formed upon mutation P436R. F) Collective variables (CVs) from MD simulations of the wild-type and P436R mutant showing changes in the catalytic residue dihedral angle D275–D462.

To explore the structural impact of S297F and P436R, we analyzed the contacts formed by S297 and P436 with their surroundings during the MD simulations of the wild-type POLE. Both mutation sites are in proximity to the catalytic residues D275 and E277. In the MD simulations of the wild-type protein, S297 forms stable atomic contacts within 5 Å with both D275 and E277 (**Figure 4B**). In contrast, P436 forms only transient contacts within 5 Å with the catalytic residue D462 (**Figure 4C**). This spatial arrangement suggests that the introduction of a bulky phenylalanine at position 297 or a charged residue at position 436 may induce conformational shifts and re-wire the intramolecular interactions in the nearby catalytic site, potentially altering its geometry (**Figure 4A**). However, it is important to note that the method used to estimate ΔΔGs does not account for backbone rearrangements, which may be relevant for these variants. We therefore performed additional one-microsecond MD simulations of the S297F and P436R variants using a construct corresponding to residues 30-522 of POLE (POLE_30-522_) to evaluate whether these amino acid substitutions induce conformational changes in the active site. We observed that the aromatic ring of the phenylalanine at position 297 in POLE-S297F_30-522_ causes a shift in the catalytic site conformation, associated with increased distances between the side chains of residues D275, E277, and D462, as well as altered reciprocal orientations (**Figure 4D**). In contrast, P436R engages in a network of salt bridges with E438 and D462, with occurrence in 67% and 90% structures from the simulation (**Figure 4E**), causing a change in the reciprocal orientation of the three catalytic residues (**Figure 4E-F**).

Supporting this hypothesis, S297F is a recurrent cancer driver mutation and has been associated with hypermutated tumor profiles [24][25]. Its damaging impact is also supported experimentally by a yeast-based functional assay [26], which links this variant to significantly elevated mutation rates, consistent with a loss-of-proofreading functional phenotype (**Figure 2A**, Experimental Data Classification row). P436R has also been experimentally verified to have a damaging effect (**Figure 2A**, Experimental Data Classification row). Specifically, mutation-rate assays in yeast showed that this mutator allele produces an elevated mutation rate relative to the wild-type in both heterozygous and homozygous contexts, although the effect is milder than observed for the S459F variant [27].

### N363K is a known pathogenic POLE variant that affects DNA interactions in both the frayed and mismatch excision states

N363K (p.Asn363Lys) is classified as likely pathogenic in ClinVar based on segregation analysis, in silico predictions and evolutionary conservation (ClinVar ID 420748). On the other hand, according to the curation of experimental data included in MAVISp, N363K is known to have a damaging effect in functional assays (**Figure 2A**, Experimental Data Classification row) [28,29]. As for other variants, we leveraged cryo-EM structures of POLE that capture different functional states during mismatch recognition and repair. Using these structures and the MAVISp protocol, we investigated the effects of N363K in conformations corresponding to different functional states (**Figure 2A**). Two states are particularly relevant for the impact of N363K, i.e., the frayed state, during which the terminal bases of the nascent DNA strand are loosely accommodated in the protein cleft, deviating from the template strand, and the mismatch-excision state, where the mismatched base is correctly positioned for removal [30]. The analyses with the local interaction module of MAVISp, which are based on binding free energies, results in N363K weakening protein-DNA binding in the frayed state while stabilizing interactions in the mismatch-excision intermediate (**Figure 2A**). As the two different functional states are sequential in the catalytic mechanism of the complex, and the frayed conformation occurs earlier than the mismatch-excision intermediate, the lower binding affinity predicted by the mutant in the frayed state could hinder the ability of even reaching the subsequent states, making the effect of the variants on them less relevant. Therefore, we focused the rest of our analysis specifically on the frayed state.

Visual inspection of the mutant and wild-type structures generated by the side-chain sampling applied to the binding free energy protocol (see Materials and Methods) shows that replacing the asparagine in position 363 with a lysine introduces additional contacts with the terminal two nucleotides at the 3′ end of the nascent strand (**Figure 5A-B**). As previously mentioned, FoldX predicts that the mutation has an overall detrimental effect on DNA binding (**Figure 2A**). Next, we evaluated the individual energy terms related to the overall change of binding energy upon amino acid substitution computed as mutant–wild-type differences for each term (**Figure 5C**). This analysis shows that the electrostatic component, likely due to new contacts introduced by the ε-amine of the lysine side chain with the phosphate group of DNA, is strengthening the interaction. Nonetheless, the dominant detrimental contribution for binding arises from a substantial difference in solvation energy. This suggests that the additional polar contacts sequester the phosphate group, hindering its ability to interact with water and thereby rendering the frayed intermediate energetically less favorable. Such destabilization might compromise formation of the prerequisite intermediate for mismatch excision, impairing proofreading efficiency. Nevertheless, the method here used for the binding free energy calculations does not include backbone flexibility and cannot capture large conformational rearrangements induced by amino acidic substitutions of the protein or the DNA. To better understand the mechanisms at play we would need to resort on molecular dynamics simulations of N363K variant in complex with DNA in the exonuclease domain, similarly to what we did above to further investigate variants effects on structural stability in the free states of POLE or POLD1.

**Figure 5.**
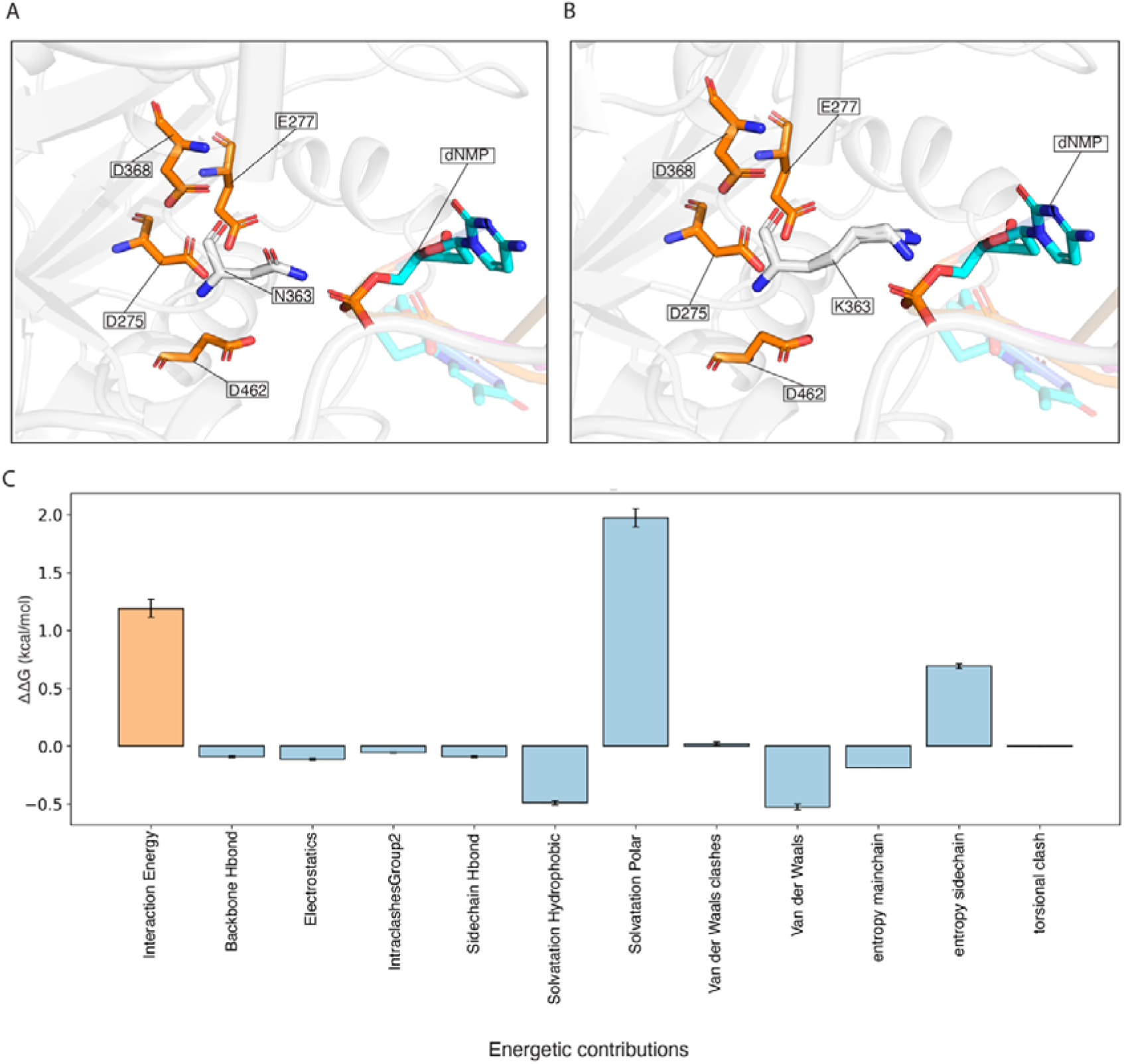
Geometrical and energetic analysis of the N363K variant and its impact on DNA binding. (A–B) Superposition of the five structural models generated by FoldX for the wild-type (A) and N363K-mutant (B) POLE in the frayed state. Exonuclease catalytic residues are shown in orange, residue 363 (Asn or Lys) in grey, and the mismatched dNMP in cyan. C) FoldX energetic decomposition of the predicted binding free energy (kcal/mol) between POLE and DNA in the frayed state for wild-type and N363K. Error bars represent standard deviations computed on the five ΔΔG obtained from the 5 independent runs of FoldX for a specific variant.

### Characterization of variants with uncertain significance or conflicting evidence in ClinVar

The comparison reported in **Table 1** allowed us to validate the default MAVISp discovery workflow for VUS and other unknown variants illustrated in **Figure 1**. The workflow relies on AlphaMissense to identify potentially damaging variants, combined with a second layer of evidence based on DeMaSk loss-of-fitness/gain-of-fitness signatures. On the variants selected in this manner, we focus on the variants that are reported with at least one identified MAVISp mechanistic indicator and explore them further for their structural alterations. Among these, we also evaluated the pLDDT scores of the original structure. In cases of low accuracy (i.e., pLDDT < 70), we verified that these variants were in flexible sites (see Materials and Methods and OSF repository) during the MD simulation. Variants with remaining low pLDDT, not explained by structural flexibility, were not considered further in the analysis of the structural mechanisms to avoid low accuracy in the predictions. Overall, MAVISp allowed us to provide structural information on 279 and 136 variants with uncertain significance or conflicting ClinVar classification of POLE and POLD1, respectively. The full set of data in the form of dotplots, lolliplots, and heatmaps is provided in the OSF repository associated with the publication for consultation (https://osf.io/z8x4j/). We then considered that, in the specific case of POLE and POLD1 and their cancer phenotype, it is more relevant to evaluate variants with a destabilizing effect on structural stability that are candidates for causing structural rearrangements of the regions important for the catalytic activity or for the coordination of the cofactors. We thus focused our analysis on the variants with a damaging prediction for stability and retained only those in the proximity of these sites (i.e., within 10 Å). We then applied the ranking protocol described in the Materials and Methods on the remaining variants (97 for POLE and 82 for POLD1). Due to the large number of annotations, we here focused on the top 25 ranking variants in detail for each protein target and applied two different approaches to highlight the most interesting predictions for further investigations (**Supplementary Figures S2 and S3**). Among these, we identified a group of potentially damaging variants that destabilize interaction with DNA, i.e., N363D, G498R, T1090R in POLE (**Figure 6A**) and G702S in POLD1 (**Figure 6B**), or with other proteins (W2276C, I1183N) in POLE (**Figure 6A)** and D1068Y, G500E in POLD1 (**Figure 6B**). Additionally, 21 variants of POLD1 and 20 variants of POLE with alterations in structural stability for further investigation. C512Y (p.Cys512Tyr) featured high in the ranking due to high predicted values of changes in folding free energy predicted by FoldX (average ΔΔG = 13.20 +/- 4.95 kcal/mol) but remained uncertain in the classification for structural stability due to mild or negligible effects predicted by RaSP (average ΔΔG = 1.73 +/- 0.50 kcal/mol). We thus verified this prediction also using the protocol for stability based on Rosetta, but the variant remained uncertain.

**Figure 6.**
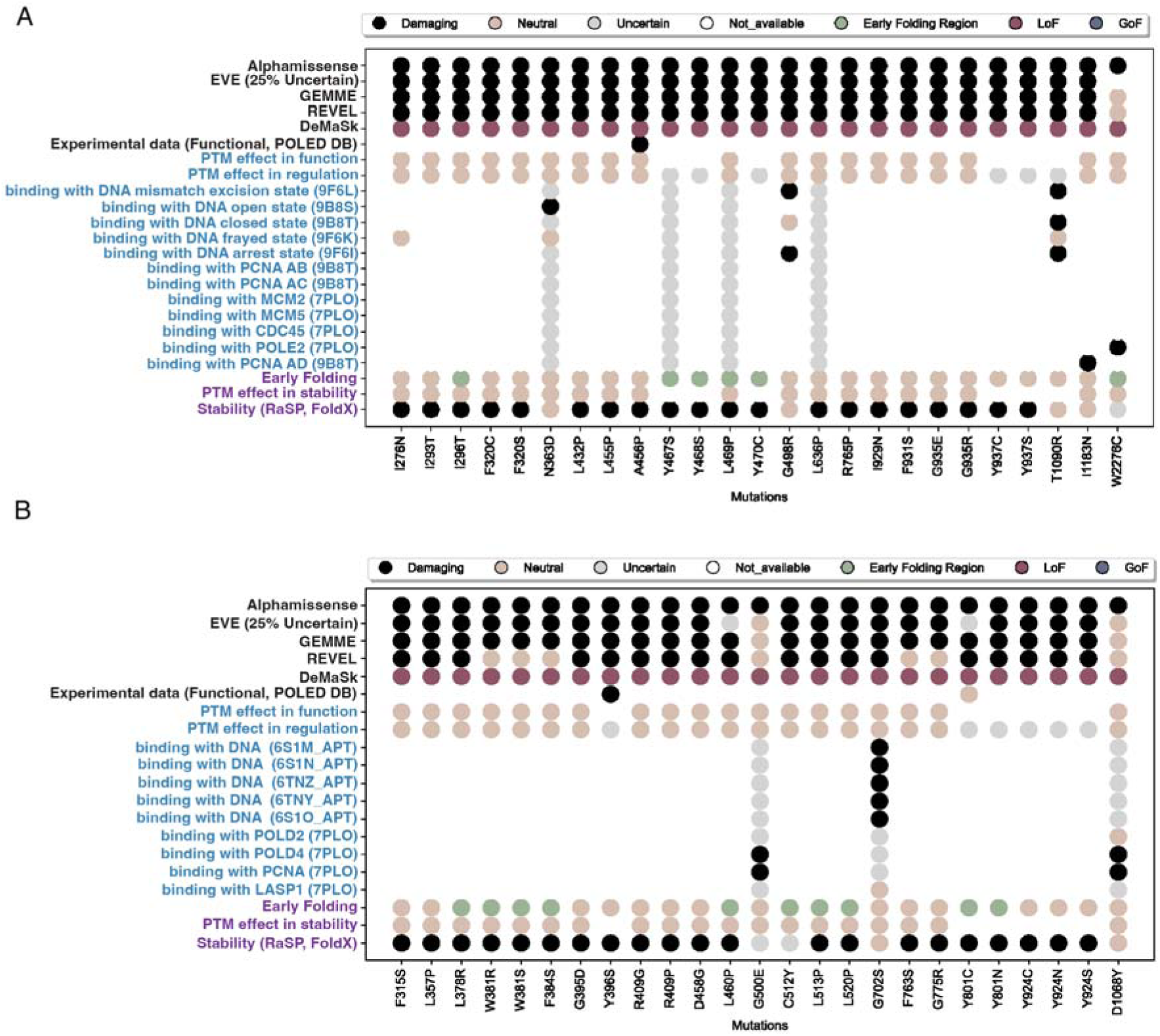
Structural features and VEP predictions for VUS and variants with conflicting evidence in ClinVar. The dotplots illustrate the different predictions and effects from the top-25 ranking VUS or variants of conflicting evidence in POLE (A) and POLD1 (B). The heatmaps with the detailed scores used in the ranking are provided in **Supplementary Figures S2 and S3**.

D1068Y (p.Asp1068Tyr) and G500E (p.Gly500Glu) in POLD1 alter both the interaction with PCNA and POLD4. Similarly, I1183N (p.Ile1183Asn) associated to hereditary cancer predisposing syndrome appears to disrupt the interaction between POLE and PCNA, Nevertheless, the destabilization of the interaction between POLD1 and its accessory subunit POLD4 and PCNA is unlikely to promote a tumorigenic phenotype, given the critical role of this interaction in ensuring replication processivity [4,31–35].

### G702S in POLD1 and N363D, G498R, T1090R in POLE are potentially damaging variants with effects on the interaction with DNA

G702S (p.Gly702Ser) in POLD1 exhibits detrimental effects on binding across all DNA-bound conformations analyzed in this study and has been previously reported in association with colorectal cancer in ClinVar (ClinVar ID 1720907). The variant lies at the interface between the fingers and palm domains, a region essential for DNA coordination and known to function as a mismatch-sensing module (**Figure 7A, left panel**). Notably, G702S is located immediately adjacent to Y701, which forms the sugar-steric gate responsible for discriminating dNTPs from rNTPs [36], [5]. To investigate the structural impact of this substitution, we examined local contacts using Arpeggio [37] visualizing the difference in contacts that the residue has with the DNA (**Materials and Methods**). The introduction of a serine side chain produces extensive van der Waals clashes with the template nucleotide complementary to the incoming substrate, as well as additional clashes with the adjacent upstream base (**Figure 7A, left panel**).

**Figure 7.**
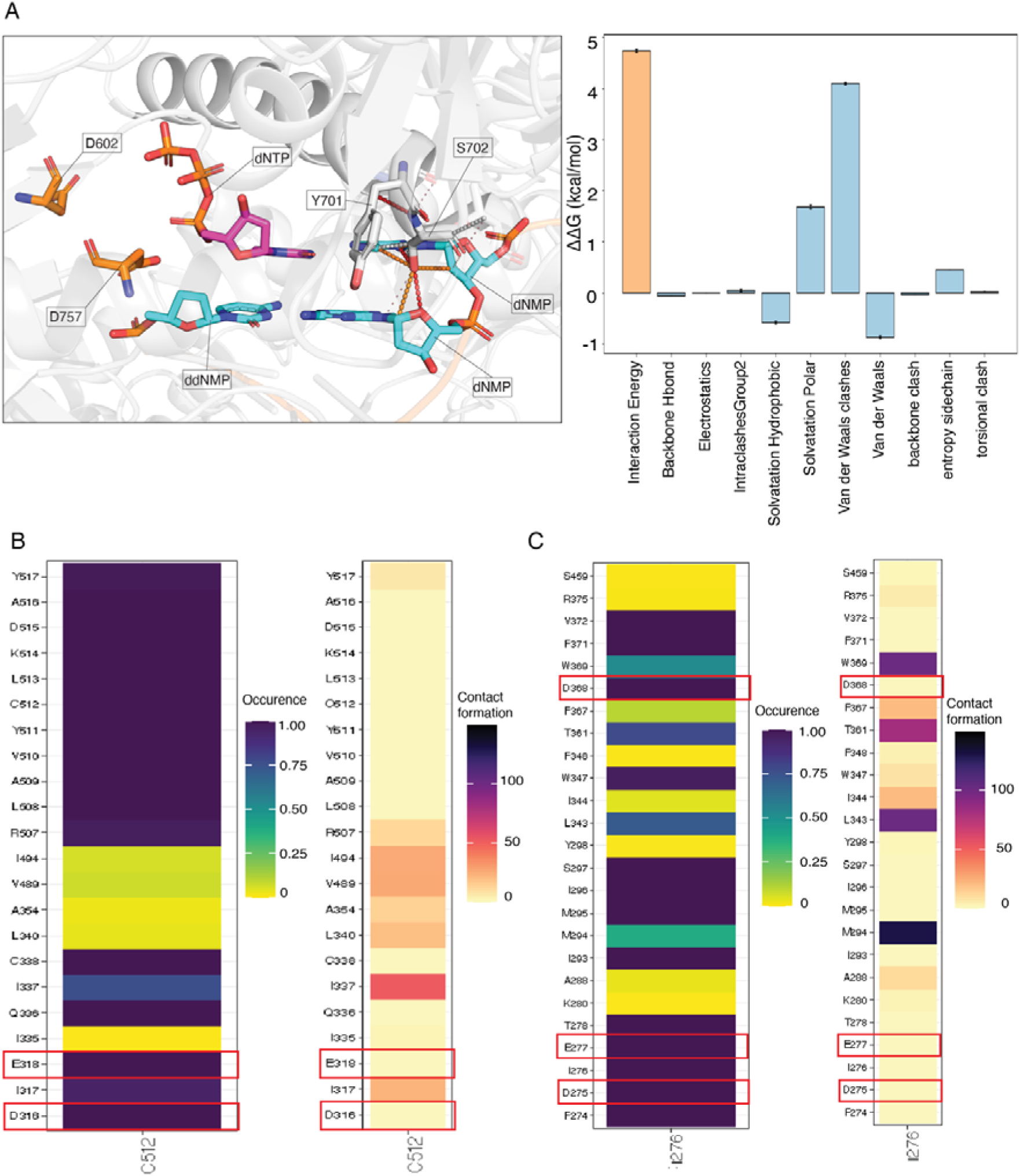
Analysis of POLE and POLD1 mutations prioritized by the MAVISp scoring framework as candidates for experimental validation. A. Left: Superposition of the five FoldX-generated structural models of the G702S POLD1 variant with the experimental POLD1 structure (PDB 6TNY), from which only the ddNMP has been shown to better represent the geometry of the catalytic site. Polymerase catalytic residues are shown in orange, Ser702 and Tyr701 in grey, the templating dNMPs in cyan, and the incoming dNTP in magenta. Protein–DNA contacts established by Ser702 are annotated according to Arpeggio interaction categories: red, van der Waals clash–polar; pink, proximal–polar; orange, van der Waals clash–weak polar; sand, proximal–weak polar; white, van der Waals clash. Right: FoldX energetic decomposition of the predicted POLD1–DNA binding free energy (kcal/mol) for the wild type and the G702S variant. Error bars indicate propagated standard deviations from five independent FoldX models generated for each state. B–C. Contact analysis for residues C512 and I276, along the POLE molecular-dynamics trajectory, illustrating the occurrence and persistence of residue–residue interactions over time (contact frequency expressed as the number of frames in which each interaction is observed). Relevant residues establishing contacts are highlighted in red.

Additionally, we analyzed the energy components of the FoldX change of binding free energy upon mutation, which shows that van der Waals repulsion predominantly drives the change in binding free energy upon mutation (**Figure 7A, right panel**). Given its position within a hotspot for nucleotide selectivity, these steric perturbations are likely to disrupt proper DNA positioning during replication and could impair polymerase activity. However, given the association of G702S with predisposition to colorectal cancer, a more plausible mechanistic consequence is a decrease in replication fidelity, driven by an increased rate of nucleotide misincorporation due to destabilized DNA interactions in a region critical for DNTP selectivity. Taken together, these features identify G702S as a possible candidate for experimental validation.

### A group of VUS in POLE and POLD1 is potentially damaging for the architecture of the catalytic site in the exonuclease domain

For the remaining variants with predicted effects on structural stability, we investigated in more detail the atomic contacts with the surrounding residues and their occurrence and strength during the MD simulation to identify cases that could behave similarly to the known pathogenic POLE variant S297F (**Figure 4**), i.e., variants that can impact the architecture of the catalytic or cofactor binding sites. Candidates in this category are C512Y (p.Cys512Tyr) of POLD1 and I296T (p.Ile296Thr) of POLE, which are in stable contacts with the catalytic residues in the exonuclease domains (D316/E318 and D275/E277, respectively) (**Figure 7B**, **Supplementary Figure S4** and OSF repository). I276N (p.Ile276Asn) variant of POLE is at a site central for contacting all three catalytic residues D275, E277 and D368 (**Figure 7C**). Additionally, F315S (p.Phe315Ser) and Y396S (p.Tyr396Ser) in POLD1 could cause conformational changes acting on D316 and D402 of the catalytic site (**Supplementary Figure S4, Figure 8A** and OSF repository). A similar effect could be exerted by POLD1 G395D variant (p.Gly395Asp), which participates only in transient interactions with D402 (OSF repository), and by the POLE L455P (p.Leu455Pro) and I293T (p.Ile293Thr) variants which engage in stable or more transient contacts with either E277 or D402 (**Supplementary Figure S4** and OSF repository). L513P (p.Leu513Pro) is also located in a site with weak interactions with D316 but has stable contacts with D515, which is expected, by homology, to coordinate the Mg^2+^ metal ion for the reaction (**Figure 8B**). All these variants could be suitable candidates for future investigation using MD simulations of the effects on the catalytic sites of POLD1, like what we performed here for the POLE S297F and P436R variants. They could compromise the proofreading activity of POLD1 or POLE. Among them, Y396S is a POLD1 VUS with a ClinVar review status of 1 and it is associated with colorectal cancer susceptibility. Supporting the prediction, a yeast-based assay using the *CAN1*, *trp1-289*, and *his7-2* reporters confirmed that Y396S leads to a high mutation rate [38] (**Figure 6B**, Experimental Data Classification row). Similar assays could provide further insight into C512Y, F315S, W381R/S, G395D, and L513P. Finally, we examined all ranked variants to identify additional candidates that may perturb either the architecture of the catalytic site (S297F-like variants) or the stability of DNA interactions within the exonuclease domain (N363K-like variants). Our analysis highlighted N363D (p.Asn363Asp), which occurs at the same position as the known pathogenic variant N363K. Similar to N363K (p.Asn363Lys), N363D (p.Asn363Asp) displays a destabilizing effect on DNA binding in the frayed state, primarily driven by unfavorable contribution in terms of backbone and side-chain hydrogen bonding, as well as solvation (**Figure 8C**). We also identified G498R (p.Gly498Arg) as affecting DNA binding in two distinct POLE conformations: the closed state, in which the polymerase is poised for nucleotide incorporation, and the arrest state, associated with detection of a misincorporated nucleotide. In both conformations, van der Waals clashes and solvation energy emerge as the dominant destabilizing components (**Figure 8C**). For this mutation, the large magnitude of the reported changes of free energy and their components in the closed state are likely connected to known limitations of FoldX itself. The large difference in volume between Glycine and Arginine, coupled with the limited sampling performed by FoldX and immovable backbone, make accommodating the replacement of the side chain more difficult for this case. Finally, we observed that T1090R (p.Thr1090Arg) perturbs both conformations acquired during polymerization (open and closed). In both the open and closed state, the detrimental effect on binding arises from changes in polar solvation effects and side-chain entropy (Figure 8C). Although T1090R (p.Thr1090Arg) lies far from the catalytic center, making a direct effect on catalytic activity unlikely, it may subtly modulate DNA binding or processivity. However, given that this variant is associated with hereditary cancer predisposition, it is unlikely that it would be positively selected if it severely compromised processivity. Further experimental investigation will be required to elucidate its precise functional impact. All candidate variants with relevant structural or energetic signatures for follow-up validation are summarized in Table 2.

**Figure 8.**
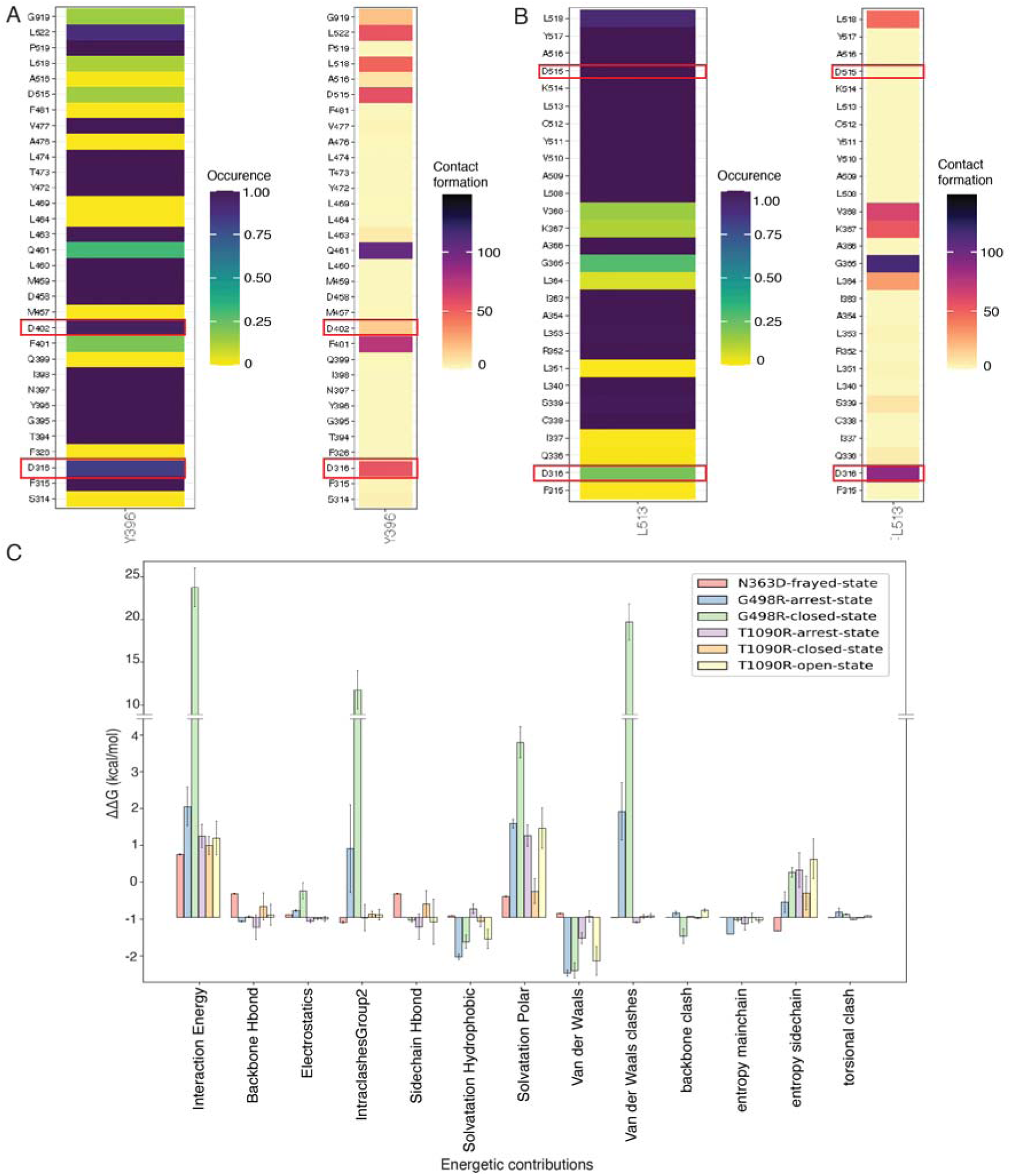
Analysis of POLE and POLD1 mutations prioritized by the MAVISp scoring framework as candidates for experimental validation. (A-B) Contact analysis for residues Y396 (A) and L513 (B) along the POLD1 molecular-dynamics trajectory, illustrating the occurrence and persistence of residue–residue interactions over time (contact frequency expressed as the number of frames in which each interaction is observed). Relevant residues establishing contacts are highlighted in red. C. FoldX energetic decomposition of the predicted POLE–DNA binding free energy (kcal/mol) for the wild type and the N363D, G498R (closed and arrested conformations), and T1090R (open, closed and arrested conformations) variants. Error bars represent propagated standard deviations calculated from five independent FoldX models generated for each state.

**Table 2.**
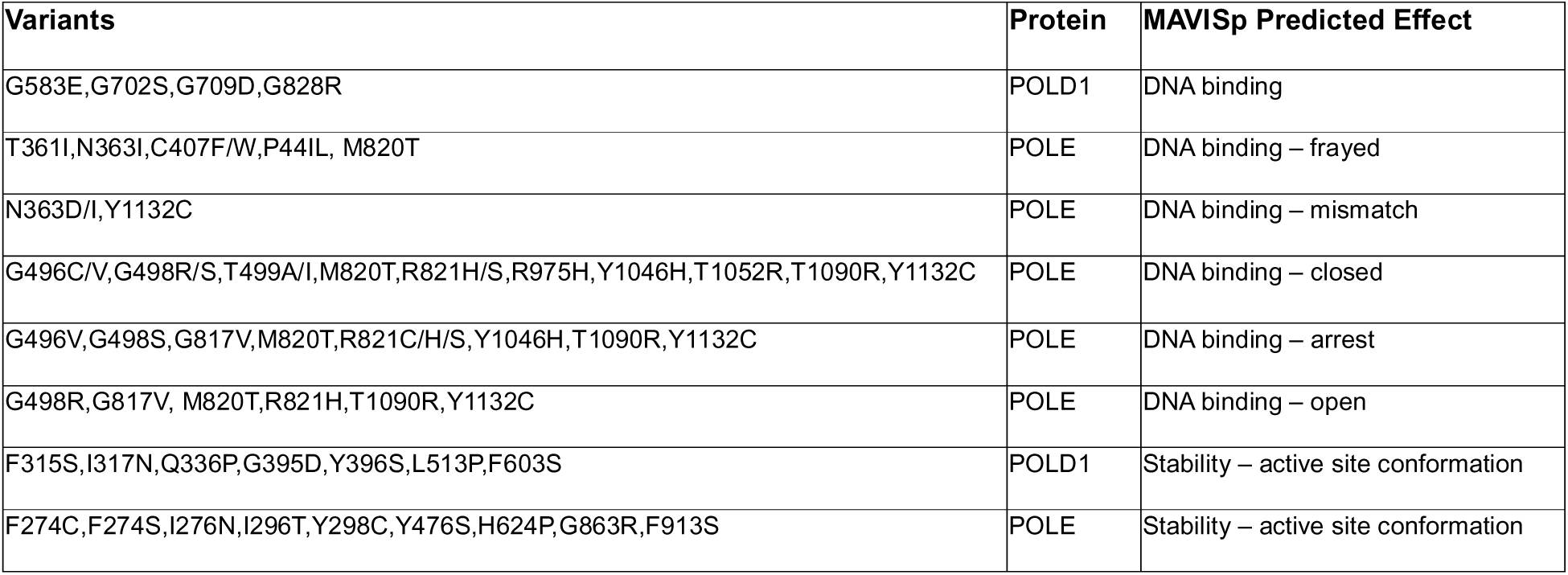
Selected VUS candidates for experimental validation. This table lists high-priority missense mutations in POLE and POLD1 for which MAVISp predicted a mechanistic impact potentially relevant in disease. The focus is on variants predicted to alter the binding interface with DNA or which could damage the structural stability and conformation of the catalytic site in the exonuclease domain or for Mg^2+^ coordination. All listed variants were predicted to be damaging by AlphaMissense and to result in loss or gain of-fitness by DeMaSk. In the case of POLD1, we reported only variants with an effect on DNA binding on at least two of the 3D structures used as representative for this interaction.

### Mechanistic indicators for potentially damaging variants reported in COSMIC and cBioPortal

MAVISp also facilitates the interpretation of variants reported in other external databases such as COSMIC and cBioPortal [39,40], offering insights into the potential molecular mechanisms underlying their functional effects. We applied the same strategy used for the VUS characterization to these classes of variants. The full set of analyses is also available in the OSF repository linked to this study (see the *downstream analyses* folder therein), applying the dot plot and lolliplot visualizations, as used for the ClinVar variants. We identified 70 and 32 variants in COSMIC and/or in cBioPortal for POLE and POLD1, respectively, with predicted damaging effect and a linked mechanistic indicator, illustrated in **Figure 9A-C**. They are related to changes in Local Interactions, PTM, or Structural Stability. S314 of POLD1 is reported as a phospho-site from studies of mass spectrometry according to PhosphoSitePlus [41] and part of a phospho-modulated short linear motif (PPI-docking motif RVXF, for recruitment of PP1 for post translational regulation). During the simulations, it remained buried from the solvent (**Supplementary Figure S5**), but it is in proximity of a loop which could undergo conformational changes not captured in the timescale simulated here, potentially exposing the site for phosphorylation. This is a common feature for cryptic phosphorylation sites [42]. The mutations to cysteine are expected to remove the possibility of this phosphorylation to regulate the structural stability in this region. A bulky phospho-group in position 314 would potentially have the effect of promoting open conformation of this region, which could alter the Mg2+ coordinating residue D515. This variant is a candidate for future studies with enhanced sampling approaches to model longer timescale dynamics and verify both the accessibility of the site and the conformational effects exerted by the phosphorylation. Additionally, many other variants that are not reported in the three databases are available with their predicted effects for exploration via the POLE and POLD1 entries in the MAVISp database (https://services.healthtech.dtu.dk/services/MAVISp-1.0/) or through the output files in the OSF repository associated with this publication (https://osf.io/z8x4j/).

**Figure 9.**
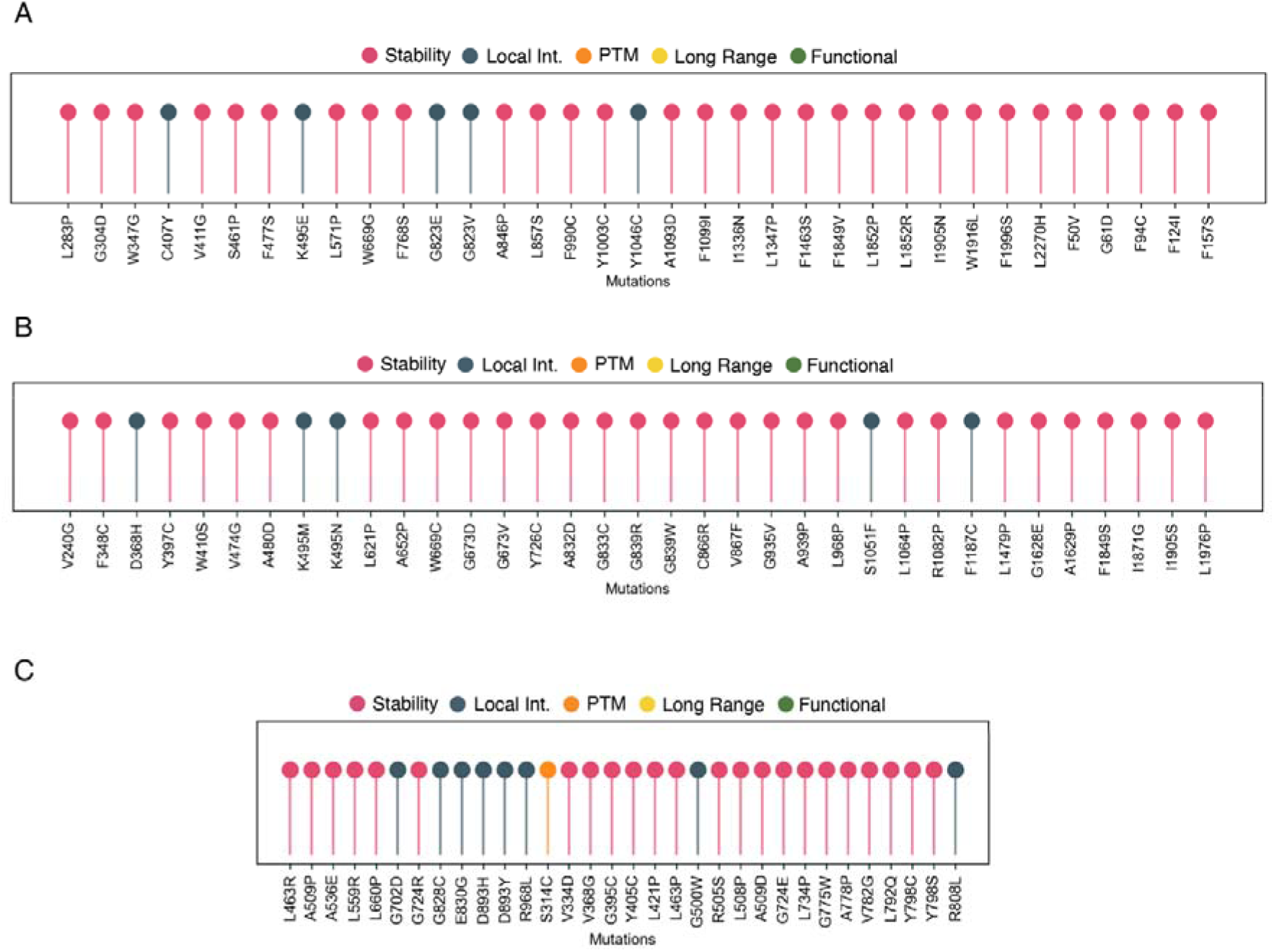
Variants of POLE and POLD1 reported in COSMIC and/or cBioPortal predicted using the default discovery workflow of MAVISp. The variants have been selected as damaging for AlphaMissense and loss/gain-of-fitness by DeMaSk. The lolliplots shows the main classes of mechanistic indicators predicted for POLE (A-B) or POLD1 (C). We can observe that most variants cause structural alterations in the proximity of the catalytic or cofactor binding sites (pink), followed by cases altering local interactions with other proteins or DNA (grey) and one case of alteration at the phosphorylation level (orange).

### Mechanistic Evidence and Reclassification Potential for ClinVar VUS

We estimated calibrated PP3/BP4 scores and structure-based PM1 supporting evidence as explained in the **Materials and Methods** for the ClinVar variants of uncertain significance (VUS) in the exonuclease domain of POLE (residues 268-471) and POLD1 (residues 304-533, **Table S2**). We restricted the analyses to this domain following recommendations from a previous work [12] according to which the exonuclease domain is the only region with sufficient known pathogenic and benign variants to allow for threshold calibration. Across 465 VUS in POLE, our integrated framework identified 212 variants that meet the criteria for PP3 Likely pathogenic candidates (**Table S3**). On the other hand, 138 met benign evidence thresholds (BP4), with 115 remaining classified as VUS in POLE. For POLD1, we evaluated 518 VUS, of which 152 were PP3 and 185 were BP4 (**Table S4)**. The remaining variants lacked sufficient computational support and remained classified as VUS. MAVISp assigned 40 and 32 variants as Likely pathogenic candidates in POLE and POLD1, respectively, with also PM1-supporting structural evidence (**Table S3-S4, Figure 10**), which were mostly linked to changes in structural stability or interactions with the DNA, and with a case of alterations for the PTM function class and one for changes in protein-protein interactions (in POLD1). Together, these findings suggests that the integration of the VEPs included in MAVISp can provide PP3 and BP4 evidence, while MAVISp predictions provide mechanistic PM1-like support for a subset of POLE/POLD1 VUS. While functional assays (PS3/BS3) remain required for formal clinical reclassification, MAVISp offers a prioritization strategy for variants most likely to have damaging structural consequences.

**Figure 10.**
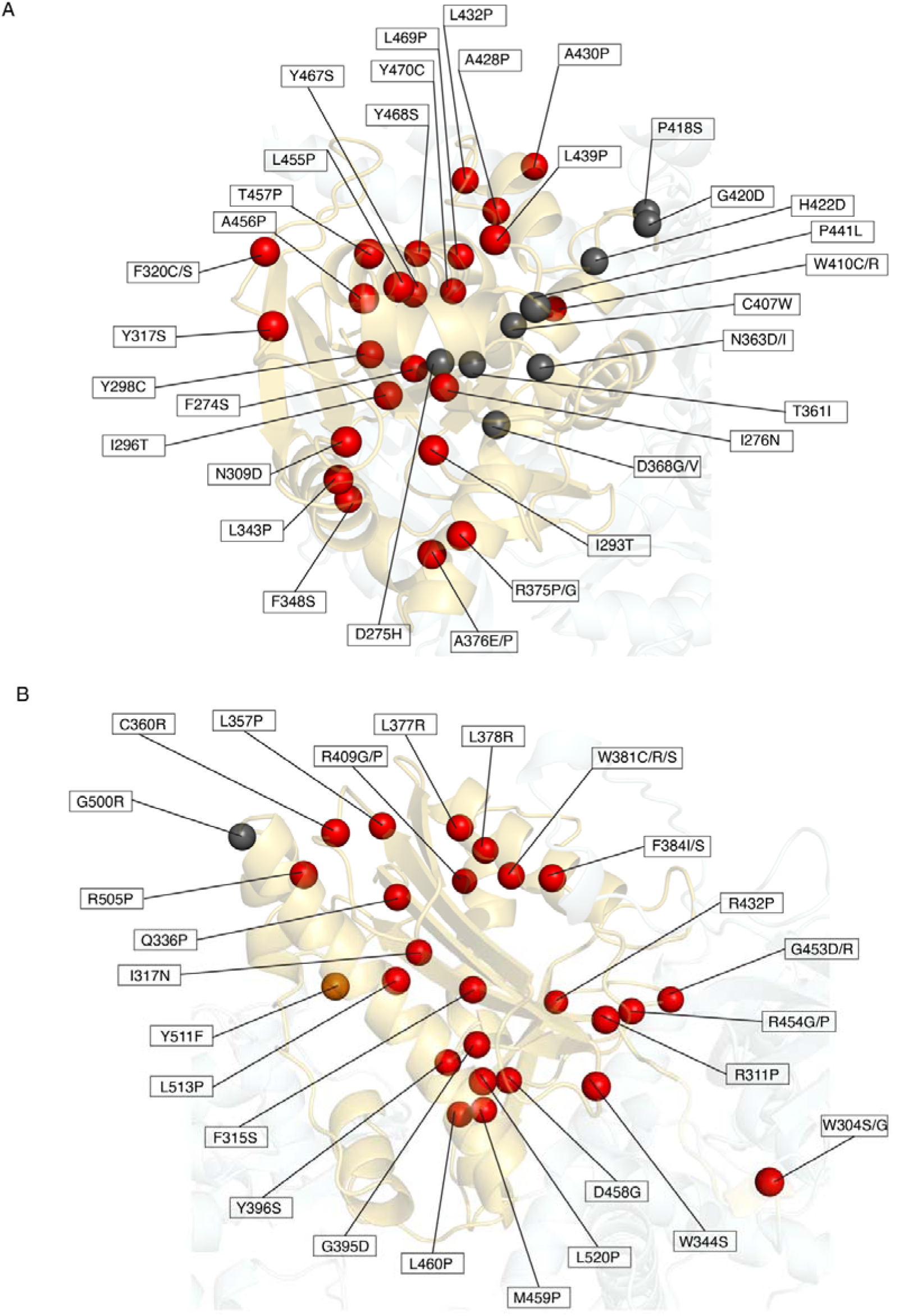
Variants classified as PP3 with additional PM1_supporting evidence from MAVISp structural mechanisms. This figure shows POLE (A) and POLD1 (B) variants that meet the ACMG PP3 criterion and additionally exhibit PM1-supporting evidence based on structural mechanisms identified by MAVISp. For each variant, the mechanistic category inferred by MAVISp is used to assign distinct colors when mapping the variants onto the protein structure (red for variants classified as damaging by the Stability module, gray for those classified as damaging by the Local Interaction module, and orange for those classified as damaging by the PTM Stability module), thereby enabling a visual comparison of the different structural mechanisms contributing to the PM1-supporting evidence.

### Conclusions

Here, we provide a comprehensive structural analysis of all possible missense variants in the DNA polymerases POLE (2-2286) and POLD1 (2-1107), using the MAVISp (Multi-layered Assessment of VarIants by Structure for proteins) framework and molecular dynamics simulations. By integrating predictions of changes in protein stability, functional sites, and alterations of DNA and protein–protein interactions, we provide mechanistic insights into both known pathogenic variants and VUS or other clinically unknown variants. A recurrent mechanism underlying the effect of the studied mutations, as identified in our analysis, is the alteration of structural stability. However, interpreting this effect in the context of cancer for these genes is particularly complex. Given the essential roles of POLE and POLD1 in DNA replication, complete protein misfolding leading to degradation is unlikely to support a tumor-promoting phenotype. Mutations that markedly reduce polymerase abundance within the cell are therefore not expected to confer a selective advantage in cancer, and are more plausibly interpreted as heterozygous passenger variants, functionally compensated by the remaining wild-type allele. Accordingly, homozygous mutations observed in cancer samples for which MAVISp predicts only a strong destabilizing effect on protein stability (ΔΔG > 3 kcal/mol), and no other mechanistic consequence, should not be automatically interpreted as causing polymerase depletion. This is particularly relevant given that ΔΔG-based predictors, including those implemented in MAVISp, model mostly local structural perturbations around the mutated residue and cannot account for global folding dynamics, especially in large, multidomain proteins such as replicative polymerases. These observations prompted us to focus on variants with predicted destabilizing effects in the proximity of the active site, and that can modulate the conformation of the catalytic residues, such as S297F and P436R, analyzed in detail in this study. On the other hand, another clear signature emerging from this investigation refers to variants that cause alteration in the DNA binding cleft, as exemplified here by N363K and G498R.

Variant interpretation must also consider the functional domain in which the mutation occurs. In both POLE and POLD1, a local destabilization of the exonuclease domain could impair the proofreading activity while preserving the polymerase function, thus contributing to a hypermutator phenotype that promotes tumor progression. Conversely, mutations in the polymerase domain require a different interpretation since the polymerase activity should not be affected. In this case, the analyses should be tailored toward the investigation of a reduction of replication fidelity. Altogether, these considerations underscore the importance of experimentally validating variants predicted to affect structural stability or DNA binding, particularly through assays assessing mutation rates and cellular proliferation. Such efforts are essential both to clarify their oncogenic potential and to benchmark the accuracy of MAVISp predictions in prioritizing clinically actionable variants.

Importantly, our framework and the downstream analysis of the molecular dynamics simulations identified several candidates for further experimental validation. The candidates can be grouped in two main categories: i) variants with damaging effects on the active site conformation and variants with damaging effects in relation to DNA recognition (**Table 2**). Additionally, our integrative approach, combining structure-based mechanistic indicators with the phenotypic context, provides a robust framework for prioritizing variants for functional validation and potential clinical reclassification. The results can guide the design of targeted functional assays, supporting both the validation of the predictions and the generation of evidence that may contribute to variant interpretation under the ACMG/AMP guidelines. In this context, we also proposed calibrated PP3 and BP4 classifications for the VUS in the exonuclease domains of POLE and POLD1, along with a subset of 73 PP3 variants with PM1-like structural evidence provided by MAVISp. In conclusion, our work highlights several structural mechanisms through which POLE and POLD1 variants may alter their function. While MAVISp captures critical functional and structural features of POLE variants, its application to POLD1 is limited by the lack of DNA-bound human structures at the different steps of the DNA repair mechanism. As new structural data emerges, especially for excision repair intermediates, the MAVISp database can be complemented with new information, to further enhance its effectiveness.

## Materials and Methods

### Identification and aggregation of variants for the study

We used Cancermuts [43] to retrieve variants from COSMIC [39] and cBioPortal [40], as well as to aggregate variants from ClinVar [7][45]. We filtered variants downloaded from ClinVar, keeping only those annotated as having a protein mutation in specific protein sequences, namely RefSeq IDs NP_006222 (POLE) and NP_002682 (POLD1). These correspond to the sequences of the main UniProt isoform. The data were collected on 24/06/2023. We applied the Cancermuts workflow (https://github.com/ELELAB/cancermuts) and a Python script clinvar.py (available in the mavisp_templates folder of https://github.com/ELELAB/MAVISp_automatization) to retrieve the variants of interest and their annotations.

### Performance of VEPs against ClinVar data

ClinVar variant interpretations from the MAVISp database files were used as the reference set to assign positive and negative labels. We used pathogenic and likely pathogenic variants as positive and benign and likely benign variants as negative. In MAVISp, a protein-level mutation can have multiple ClinVar classifications, as multiple variants at genome level can result in the same protein-level variant. For these cases, a variant was considered positive if all the available variant interpretations were pathogenic or likely pathogenic, negative if all the available variant interpretations were benign or likely benign, and excluded from the dataset for cases of mixed intepretation (e.g., “pathogenic/benign”) or if not all interpretations allowed us to classify them (e.g., “uncertain significance”, “conflicting interpretations”). Variants without a resolvable positive/negative assignment were removed from performance analyses. ClinVar review status was parsed directly from the corresponding column, which in the MAVISp output is provided as a numeric value (0–4 stars). Performances were assessed by comparing each predictor binary classification output to the ClinVar-derived ground-truth labels. Only variants for which all predictors produced a valid classification were included in the final evaluation set. For each method, we calculated Recall, Specificity, Accuracy, Matthews Correlation Coefficient (MCC), F1-score, and Balanced Accuracy. MCC is robust to class imbalance and was therefore used as the primary performance indicator. The binary classifications for AlphaMissense and EVE were directly taken from the MAVISp csv file, whereas for GEMME and REVEL, we implemented further analyses to process the GEMME and REVEL scores reported in the csv file and applied different thresholds suggested by our previous studies or other publications to evaluate how they performed on the specific case study of POLE and POLD1. In details, for REVEL, we tested the performances applying a threshold of 0.5 and 0.7 [18,44], whereas for GEMME, we compared the thresholds derived by the benchmark in the MAVISp publication [8] of <= −3 for damaging variants, with a recent classification based on a Gaussian Mixture Model (GMM). The GMM divides the variants into mild, neutral, and impactful. In the comparison, we considered predictions of variants as impactful or mild as positive predictions.

### Retrieval of structure from the Protein Data Bank and the AlphaFold Protein Structure Database

We applied the STRUCTURE_SELECTION module of MAVISp [8] to retrieve the AlphaFold (AF) models from the AlphaFold Protein Structure Database [45], along with information on structures available in the Protein Data Bank (PDB) [46]. We compared the quality of the AF models against the available experimental structures of both POLE and POLD1 and ultimately selected the catalytic subunits of both proteins (POLE_2-2286_ and POLD1_2-1107_) in the AF models for our analyses. Zinc ions were modelled on the structures by superimposition to an X-ray structure (PDB ID 5VBN, chain A [47] and a Cryo-EM structure (PDB ID 6TNY, chain A [48]), to POLE and POLD1, respectively. The sulfur-iron cluster (SF4) in the C-terminal CysB motif of POLD1, which is required for successful DNA synthesis and repair mediated by the enzyme, and whose loss implies a slight destabilization of the protein [49], has not been included in the structure. This is due to its inability to fit its structure in the AF model, so that it would be correctly bound to the four conserved cysteine residues (Cys1058, Cys1061, Cys1071, Cys1076) that coordinate it, as well as the lack of parameters in the tools used in the MAVISp *simple mode* to represent this cofactor.

### Starting structures for analyses of protein complexes

Concerning the selection of structures to be used in the LOCAL INTERACTION module, we first identified interactors for POLE and POLD1 by mining the Mentha database [50], using the Mentha2PDB script from the MAVISp toolkit (https://github.com/ELELAB/PPI2PDB), and focused on interactors with a high Mentha score and available structural data. In the case of POLE, we used a Cryo-EM structure with 2.80 Å resolution (PDB ID 7PLO; [51]), in which POLE (chain A) interacts with POLE2 (chain B; POLE_1371-2280_-POLE2_1-527_), MCM2 (chain D; POLE_1371-2280_-MCM2_169-893_), MCM5 (chain E; POLE_1371-2280_-MCM5_21-651_), and CDC45 (chain C; POLE_1371-2280_-CDC45_1-566_) and a cryo-EM structure with 2.95 Å resolution for the POLE-PCNA complex (PDB ID 9B8T [52]). We could not include the Zn^2+^ ion coordinated by Cys2158, Cys2161, Cys2187, and Cys2190 in the model by superimposition, as it would have created steric clashes even after optimization of the rotamers of the surrounding residues. Regarding POLD1, we selected a Cryo-EM structure of resolution 3 Å (PDB ID 6TNY [48] to study the interaction between POLD1 (chain A) and POLD2 (chain B) (POLD1_78-1107_-POLD2_1-462_). We excluded the iron-sulfur cluster from the C-terminal domain of POLD1, due to the same limitations we encountered for POLE. As the interfaces between POLD1 and POLD4 or PCNA included a few missing residues, we modelled the complex using AlphaFold-multimer and selected the top-ranking model after a quality control step based on the Predicted Aligned Error (PAE) score of AlphaFold and the pDOCKQ2 score (>0.23) [53]. For POLD1, we identified two pre-calculated and experimentally validated models [54] with pDOCKQ2 scores > 0.3, POLD1-C9ORF169 and POLD1-LASP1. Of these, only POLD1-LASP1 was included in the LOCAL_INTERACTION analyses, having discarded the POLD1-C9ORF169 model due to its high level of disorder. Additionally, we generated models for all the remaining interactors reported by Mentha2PDB using AlphaFold3 (https://alphafoldserver.com/) and retained for the LOCAL_INTERACTION only the models that passed the quality control in terms of different scores (as explained in the original publication of MAVISp [54]. No additional structure passed the quality control steps with respect to the ones described above.

### Starting structures for analyses of protein-DNA complexes

We retrieved the structure of POLD1 in complex with DNA from a Cryo-EM structure (POLD1_78-1107_-DNA) with a resolution of 3.08 Å (PDB ID 6TNY, chains A, P, and T [48]). We retrieved the structures of POLE bound to DNA within the polymerase domain in both the open (PDB ID 9B8S, residues 27-1198, chains A, P, and T [52]) and closed (PDB ID 9B8T, residues 24-1198, chains A, P, and T states [52]). The corresponding PDB files were processed to remove molecules not supported by FoldX [55]. Specifically, the iron-sulfur cluster was removed from both structures, and the dTTP residue representing the entering dNTP was excluded from the 9B8T structure. Additionally, the A275 and A277 residues were retro-mutated to D275 and E277, respectively, using the *mutagenesis wizard* function of PyMOL [56], selecting optimal rotamers based on clash and steric hindrance considerations. In addition, four structures were employed to study the impact of mutations on the binding of POLE’s exonuclease domain to DNA in various states: post-insertion (PDB ID 9F6I, residues 27-1175, chains A, P and T, with a resolution 3.30 Å, [30], arrest (PDB ID 9F6J, residues 27-1175, chains A, P and T, with a resolution of 3.90 Å [30]), frayed substrate (PDB ID 9F6K, residues 27-1175, chains A, P and T, with a resolution of 4.20 Å [30]), and mismatch excision (PDB ID 9F6L, residues 27-1175, chains A, P and T, with a of resolution of 3.90 Å [30]). All PDB files were processed to remove the 4Fe4S clusters. In the 9F6I structure, the substrate analog DDS and a Ca²⁺ ion were removed, and, in both 9F6I and 9F6J, the A275 and A277 residues were retro-mutated to D275 and E277.

### Annotations from experiments

We included in the EXPERIMENTAL DATA module of MAVISp, the experimental results reported in the PolED database (https://poled-db.org), which aggregates literature-based data on the functional impact of POLE and POLD1 variants, including biochemical properties, cellular activity, and other phenotypic effects. The classifications reported in MAVISp generally reflect the phenotypes observed in studies that are predominantly tumor-focused, emphasizing increased mutation rates or loss of replication fidelity. Accordingly, variants are labeled as ‘damaging’ or ‘neutral’ in the final MAVISp output. ‘damaging’ corresponds to variants associated with significant defects in replication fidelity or hypermutagenic behavior. The more detailed metadata for consultation are provided in the OSF repository associated with this publication.

### Metal ion binding sites

Although we model the zinc metal ions at the metal binding sites of POLE and POLD1, some of the predictors used do not account for the presence of heteroatoms and would discard them from the analysis. This is likely to affect the free energy calculations for binding and stability at these sites. We thus applied a quality control on the csv file to separate the variants in residues coordinating the metal ions. For these variants, we did not consider the consensus approach for STABILITY or LOCAL_INTERACTIONS since RaSP or Rosetta do not include the heteroatoms in their calculations. FoldX allows the inclusion of the zinc metal ions, but it might not properly predict their physico-chemical properties. Overall, the results on stability at mutation sites in direct contact with zinc or with the sulfur/iron cluster of POLD1 have not been considered in the analysis of the selected mechanistic indicators.

### Molecular dynamics simulations

The AF models of the catalytic subunits of POLE (POLE_2-2286_), POLD1 (POLD1_2-1107_) were used as starting structures for 500-ns molecular dynamics (MD) simulations (1000 ns for the POLE_30-522_ P436R variant). In addition, we included MD simulations of a shorter POLE construct (POLE_30-522_) and its mutated variants (S297F and P436R). MD simulations were performed using GROMACS 2022 or 2024 [57] in the NVT ensemble at 298 K. Histidine residues were protonated following the assignment provided by GROMACS, except for H404 in POLE, which was manually protonated on Nδ (HSD). We carried out all-atom explicit-solvent MD simulations of POLE and POLD1 in their free state (i.e., DNA-unbound) using the CHARMM36m force field [58]. We used a dodecahedral box of TIP3P water molecules with a minimum distance between the protein and the box edges of 15 Å, applying periodic boundary conditions. We used a concentration of NaCl of 150 mM, neutralizing the net charge of the system. Bond lengths involving hydrogen atoms were constrained using the Linear Constraint Solver (LINCS) algorithm [59]. Non-bonded interactions were handled using a 0.9 nm cut-off for both van der Waals and electrostatics, with long-range electrostatics treated using the particle-mesh Ewald (PME) method [60] [61]. The system was equilibrated in multiple steps. Initially, temperature equilibration was performed under NVT conditions with position restraints applied to the heavy atoms of the protein. This was followed by pressure equilibration in the NPT ensemble in two steps: first with restraints on the protein backbone, and then without any restraints. In all simulations, temperature was kept constant at 298 K using the velocity-rescale thermostat, and pressure was coupled to 1 bar using either the Berendsen [62] or Parrinello-Rahman [63] barostat. We collected 500 ns of MD simulation for each system, according to the default MAVISp protocol, when systems contain more than 100,000 atoms upon solvation. The input, parameters, and trajectory files are reported in the OSF repository (https://osf.io/z8x4j) for each protein.

### χ1 dihedral analysis from MD simulations

In the MAVISp STABILITY module in the *ensemble mode,* the folding free energy calculations are carried out on structures extracted from the simulation. This is done to mitigate the limitations due to backbone flexibility during FoldX and RaSP/Rosetta calculations. As a default, MAVISp uses 25 frames extracted from the trajectories at regular intervals for MutateX and the structures of the three main populated structural clusters of the simulation for Rosetta or its machine learning derivative, RaSP (see original MAVISp publication for more details, [64]). This technical choice was originally dictated by the fact that the Rosetta *cartesian* protocol was computationally intensive for many structures and a saturation mutagenesis approach. Whereas RaSP does not suffer from such limitations. Thus, in focused studies on specific protein targets, as the one here presented, before using the results for further analyses or interpretation of variant effects, we apply a protocol based on comparisons of dihedral conformational states of the mutation sites to evaluate if the usage of only three main populated clusters is sufficient to recapitulate the properties observed by the entire MD trajectory. We focused on the side-chain χ1 dihedral as a probe of the sampling quality, since the free energy calculation scheme applied by the STABILITY module is highly reliant on rotameric states of side chains in the conformational sampling that they perform to derive the changes in free energies upon amino acid substitution. To this goal, we extracted with the GROMACS toolkit the χ1 torsion angles for each mutation site (excluding glycine and alanine residues). To assess whether the selected structures for the STABILITY ensemble mode were sufficient to reproduce the conformational behavior observed in the full reference trajectory, we computed the Jensen–Shannon divergence (JSD) between the probability distributions of χ1 sampled in the reference MD simulation and those obtained from each structural ensemble used to calculate free energy changes in MAVISp. Probabilities densities were estimated using normalized histograms (60 bins over −180° to +180°). This procedure has been performed independently for each residue of the protein, obtaining one DJS value for residue. Rotamer populations were additionally calculated for reporting purposes using a standardized classification: g⁺ (0–120°), trans (120–180° and from −120 to −180°), and g⁻ (from 0 to −120°). A residue was considered conformationally divergent between ensembles when the Jensen–Shannon divergence exceeded a user-defined threshold (typically JSD ≥ 0.3–0.4), chosen to allow minor sampling-related variation in peak heights while detecting substantial differences in the underlying rotameric landscape. In brief, we applied this strategy because the reference trajectory typically contains many more frames than the subsampled ensembles, so moderate differences in peak heights are expected due to sampling noise. Therefore, we used a tolerant JSD threshold (JSD ≥ 0.3–0.4) to detect only substantial conformational differences, while allowing minor deviations in rotamer populations that nonetheless preserve the correct locations of the g⁺, trans, and g⁻ peaks. The full set of results is reported on OSF. We observed that extracting 25 frames from the simulation at regular intervals resulted in JSD values above 0.4 for only a minority of sites. We thus relied on the data from RaSP and FoldX calculations using 25 frames for the variant effect analyses discussed in the Results section. Furthermore, the protocol here described was also used for selected mutation sites to better explore the agreement between the conformation used in the free energy calculations and the MD trajectory in terms of dihedral states of the surrounding of a mutation site (e.g., S297 or P436 in POLE).

### Analysis of salt-bridges

We used PyInteraph2 [65] to estimate the salt bridges formed by P436R and their occurrence in the MD simulation of this variant. We used a distance cutoff of 5.0 Å between the charged groups of the arginine at position 436 and glutamate or aspartate residues.

### Evaluation of site flexibility based on Root-Mean-Square Fluctuations from MD simulations

To quantify local backbone flexibility across the protein structure, we analysed the average C-alpha root-mean-square fluctuation (RMSF) of each residue from the (MD) simulations using GROMACS. The averages were calculated every 100 ns from the MD simulations. We remove the first three N- and last three C-terminal residues from the analyses due to the risk of artificially elevated flexibility for the classification based on RMSF. The remaining residues were used to define a reference distribution representing the protein flexibility profile. To classify the flexibility of individual mutation sites, we quantified how much the RMSF value at each site deviated from the overall flexibility pattern of the protein. First, RMSF values were calculated for every residue by averaging across the simulation, as explained above. Then, to determine whether a given site was rigid or flexible, we compared its RMSF value to the distribution of RMSF values observed for the entire protein (excluding the terminals). We assessed deviations using Z-scores, which provide a normalized way of expressing how far a value lies from the typical behavior of the system. In this context, a Z-score represents the number of standard deviations that a residue RMSF value lies above or below the mean RMSF of all the residues considered. Thus, a residue with a Z-score of zero shows the same flexibility as the average internal residues, positive Z-scores correspond to more flexible sites, while negative reflect more rigid ones. Based on this measure, we labelled each mutation site as: i) ‘rigid’ if its RMSF value fell approximately one standard deviation below the average RMSF, ii) ‘flexible’ if its RMSF value was about one standard deviation above the average RMSF, and iii) ‘partially flexible’ if its RMSF value was close to the internal average and did not deviate strongly in either directions.

### Effects of variants on protein structural stability

To estimate the changes in folding free energy upon mutation (ΔΔGs), we used the STABILITY module in *simple mode* of MAVISp [8] using the AF models of the catalytic subunits of POLE (POLE_2-2286_) and POLD1 (POLD1_2-1107_). We therefore applied both MutateX [23], which employs FoldX 5 [55], as well as RosettaDDGPrediction, which utilizes the Rosetta cartddg2020 protocol [66] with the ref2105 energy function [67]. In addition, we applied RaSP [64], a machine learning model trained on Rosetta. While Rosetta was only applied to a selection of variants due to limitations in available computational power, both MutateX and RaSP were employed for performing saturation mutagenesis scans. We used the workflows of the MAVISp toolkit for each of these methods (https://github.com/ELELAB/mutatex, https://github.com/ELELAB/RosettaDDGPrediction, and https://github.com/ELELAB/RaSP_workflow). According to the MAVISp framework, variants were classified as destabilizing if consensus ΔΔG predictions from MutateX and RaSP or from MutateX and Rosetta exceeded 3 kcal/mol, neutral if ΔΔG predictions ranged between −2 and 2 kcal/mol, and as stabilizing if both predictions agreed on ΔΔG ≤ −3 kcal/mol. Furthermore, we labelled the predicted effect as uncertain if any of either values fell between −3 < ΔΔG < −2 kcal/mol, or 2 < ΔΔG < 3 kcal/mol, or in case of disagreement between the methods.

Additionally, in specific cases of interest, we used the self-scan function of MutateX [23] with and without the repair step of FoldX as a control experiment to evaluate the quality of the initial structures before and after the repair. This approach consisted of mutating each residue to its own type. The ΔΔG values associated with self-mutations are expected to be as close as possible to 0 kcal/mol. Any deviations from this value in the repaired model (i.e., repair function active) or the initial structure (i.e., repair function inactive) indicated an initial unfavorable orientation of the sidechain and/or the surrounding residues. The self-scans, with the repair function on and off, confirmed the quality of the initial structures. No ΔΔG values were found above or below the cutoffs of ±3 kcal/mol in either the X-ray structures or in the AF models.

### Effect of mutations on early folding events

We employed the EFOLDMINE module of MAVISp, which predicts whether a specific mutation site constitutes a seed of an early folding event based on EfoldMine [68]. We applied a threshold of 0.169 for the EfoldMine score to define sites involved in early folding events and considered regions with a minimum length of three early folding residues to avoid isolated peaks [68].

### Effects of mutations on binding free energies

To estimate the changes in binding free energies upon mutation, we applied the LOCAL_INTERACTION module of MAVISp [8] based on FoldX 5 [55] and the RosettaDDGPrediction flexddg protocol [66][69] with the talaris2014 energy function [70] [71] [72]. The calculations were carried out for each complex (see OSF repository), retaining only mutations within 10 Å of the protein-protein interface to accommodate the local sampling carried out by both FoldX and Rosetta. The MutateX [23] and RosettaDDGPrediction workflows used for the data collection are available and maintained on GitHub (https://github.com/ELELAB/mutatex and https://github.com/ELELAB/RosettaDDGPrediction). After collecting predicted changes of free energy using both methods, we classified the variants according to their ability to influence binding, using a consensus scheme similarly to what done for the STABILITY module. Variants were initially classified according to each method independently, considering ΔΔG < −1 kcal/mol as Stabilizing, −1 <= ΔΔG <=1 as Neutral, and ΔΔG > 1 as Destabilizing. For any mutation, if the two methods agree on the classification (i.e. they classify the variant in the same way), then that is the final classification for the mutation; if not, the mutation is classified as Uncertain. Finally, if the estimation of ΔΔG is not available for any given mutation, it still classified as Uncertain if its side-chain relative accessible surface area (as calculated by NACCESS) is >= 25% and not classified otherwise.

### Analyses of interactions between protein and DNA

Amino acid substitutions located within the exonuclease or polymerase domains predicted to affect protein–DNA interactions were further analysed from both energetic and geometric perspectives. Energetic effects were assessed using FoldX, while geometric alterations were evaluated in terms of mutation-induced changes in interatomic contacts using the Arpeggio software (on the conformations generated by FoldX).

The analysis was carried out using a dedicated computational pipeline that integrates the extraction, aggregation, and visualization of FoldX and Arpeggio outputs (https://github.com/ELELAB/mavisp_accessory_tools/tree/main/tools/FoldX-Arpeggio_analysis). Following the MutateX execution for each variant, the individual energetic terms contributing to the final FoldX ΔΔG were extracted from each of the five structural models generated by FoldX for both the wild-type and mutant states. For each energetic component, mutant–wild-type differences were computed independently across the five models. Mean values and standard deviations were calculated from the five wild-type and five mutant models generated for each variant. Subsequently, the averaged difference between mutant and wild-type states was derived in order to identify the energetic components primarily driving the overall ΔΔG predicted by FoldX. Atomic contacts between protein and DNA were computed on individual structural models using a custom Python workflow built around the Arpeggio software suite [37] https://github.com/ELELAB/mavisp_accessory_tools). Parameters for Arpeggio were provided through a YAML configuration file specifying the distance cutoff, van der Waals compensation factor, and pH (default values: 5 Å, 0.1, and 7.4, respectively). Hydrogen atoms were not added by default and thus not locally minimized. Sequence-adjacent interactions and ambiguous contact types were excluded based on configuration-file settings. Additional details on Arpeggio are available through the project’s web interface (https://biosig.lab.uq.edu.au/arpeggioweb) as well as in the GitHub repository hosting the Python implementation used in this study (https://github.com/PDBeurope/arpeggio). The analysis has been performed on the five mutant models generated by FoldX, using the repaired wild-type structure (FoldX RepairPDB) as reference. For each model, the Arpeggio results were aggregated, and the atomic contacts and steric clashes that were gained or lost upon mutation were systematically extracted.

### Effects of mutations at phosphorylation sites

We applied the MAVISp PTM module to predict the effects of mutations at known phosphorylatable sites on the protein [8]. Briefly, the module uses multiple sources of information and custom decision logic to predict whether each mutation could have consequences on regulation by phosphorylation (PTM_regulation), structural stability (PTM_stability), or function (PTM_function) due to the interplay between the abolished phospho-site and the effect of the amino acid substitution. The data sources used to infer this includes the solvent accessibility of the mutation site calculated using NACCESS [73] on the protein structure, whether the mutation is part of a phosphorylatable linear motif as determined by ELM [74] using Cancermuts [43], as well as, the predicted changes of stability or binding free energies as calculated by FoldX 5 [55]. In the interpretation of the results from the PTM modules, as the PTM_regulation module integrates proteomics data with structural analysis of solvent-accessible surface area (SASA) to estimate the likelihood of phosphorylation at a given site, it should be applied as an initial filter to exclude residues that are sufficiently buried and thus unlikely to be accessible to kinases. In cases where the effect at this layer is predicted to be neutral or uncertain, the results from the other two modules should be disregarded.

### Effects of mutations on long-range communication to functional sites

We applied the MAVISp LONG_RANGE module to predict the mutations with allosteric effects in both the *simple* and *ensemble* modes, following the revised and benchmarked protocol recently published (https://doi.org/10.1016/j.jmb.2025.169359). Briefly, the module is based on the coarse-grained model implemented in AlloSigMA2 [75], which simulates the effects of perturbations across the protein structure. The model represents mutations as either UP or DOWN, relating to the change in side-chain size (increase; UP and decrease; DOWN), upon mutation, with respect to the wild-type residue.

In the MAVIS *simple mode*, the module applies a systematic filtering workflow to identify candidate mutations with allosteric effects (https://github.com/ELELAB/MAVISp_allosigma2_workflow). Namely, for each mutation that resulted in a significant change in residue size (at least 5 Å^3^), we identified all the predicted response residues in the protein that displayed a significant positive or negative change in allosteric free energy (|Δg| > 2 kcal/mol). We further filtered this list of residues by removing those in direct atomic contact (≤ 5.5 Å), to capture only genuine distal effects. Furthermore, the response residues were further filtered to retain only those solvent accessible (relative side chain solvent accessible surface as calculated by NACCESS [73] >= 25%) and predicted to be localized within pockets by FPocket [76,77]. Finally, the *simple* mode module classifies each mutation as stabilizing if it solely caused negative changes in allosteric free energy, destabilizing if the changes were positive, mixed effects if both changes were observed in both directions, and neutral if no significant change was predicted. Mutations that did not result in significant changes in residue side-chain volume were classified as uncertain.

In the *ensemble* mode, the LONG_RANGE module employs graph-based methodologies on the MD trajectory to refine and validate predictions of the *simple* mode module. The module uses atomic contact-based protein structure networks (acPSN), initially proposed by Vishveshwara’s group [77], constructed using the PyInteraph2 framework [78] for analysis of molecular dynamics (MD) simulations, as detailed in [79]. To define the edges of the network, we considered residue pairs with a sequence separation greater than one and a distance between their atoms of less than 4.5 Å. Residue-specific normalization factors were applied to account for their inherent propensity to form contacts. The interaction strength between two residues was calculated by dividing the number of atom pairs within 4.5 Å by the square root of the product of the normalization factors for each residue, then multiplying by 100 [77]. If the interaction strength exceeded a predefined threshold (I_min_ = 2.5 for our simulations), an edge was created, weighted by the calculated strength. Furthermore, only edges present in more than 50% of the frames across the ensemble were retained in the final acPSN. The acPSN is then weighted according to the average interaction strength. Subsequently, we then utilized the *path*_*analysis* tool of PyInteraph2 [65] to identify whether long-range communication routes exist between each identified mutation site and the respective response sites previously identified by the AlloSigMa2 workflow in the *simple* mode predictions. Shortest paths of communication are calculated, utilizing the Dijkstra algorithm [80] as implemented in NetworkX [81]. The workflow then filters the identified paths, retaining only those that span four or more residues in length, to ensure only long-range communication paths are considered. We generated the data using the code available for the LONG_RANGE data collection in the *ensemble* mode at https://github.com/ELELAB/MAVISp_allosigma2_workflow. The module then classifies the given mutation as having a damaging long-range effect if a path is identified for at least one the predicted response sites, serving as a validation of the *simple* mode results.

### Effect of mutations on active and cofactor binding sites

We employed the MAVISp FUNCTIONAL_SITES module to predict changes in the local inter-residue contacts due mutations occurring at residues within the second coordination sphere in relation to active site residues, or residues binding cofactors. Results obtained in this way are designed to understand whether side-chain replacement significantly changes the contacts of residues at or around functional sites, which might have an impact on activity or cofactor binding. To perform the contact analysis, we used Arpeggio [37] on the POLE and POLD1 wild-type structures, as well as POLE_30-522_ S297F protein structures [83][84]. The analysis has been performed using a van der Waals cutoff of 0.1, an interacting cutoff of 5.0 Å, and a physiological pH of 7.4. Subsequently, the output was preprocessed to exclude clashing contacts, attributed to poorly modelled residues, as well as contacts classified as “proximal,” which the Arpeggio [37] framework deems non-critical (https://github.com/ELELAB/mavisp_accessory_tools). In the MAVISp database, we annotated as damaging in functional sites those variants that were found to be in contact with residues of functional relevance as detailed here, or as neutral otherwise.

### Analysis of variants affecting splicing sites

Before starting to evaluate the mechanistic indicators with structural methods for selected variants from the MAVISp discovery workflow, we verified that they were not affecting splicing sites (**Tables S5 and S6**). To this goal, we evaluated the potential impact of each mutation on splice sites using SpliceAI [82] and Pangolin software [83]. Specifically, we employed the SpliceAI_lookup API from the Broad Institute (https://spliceailookup.broadinstitute.org) and used a Python script to automate the collection of results (https://github.com/ELELAB/mavisp_accessory_tools). In detail, we predicted the effects of genomic mutations within a 500-nucleotide window upstream and downstream of the mutation site. Using SpliceAI [82], we obtained predictions on whether a specific nucleotide gains or loses splicing acceptor activity, as well as whether it gains or loses splicing donor activity. With Pangolin [83], we assessed whether a specific nucleotide acquires or loses splicing activity. Each prediction is based on scores representing the difference between the reference score (in the absence of mutation) and the alternate (ALT) scores (in the presence of mutation) obtained from SpliceAI and Pangolin for a given position. These scores reflect the likelihood of the position becoming an acceptor or donor site or losing its acceptor or donor activity (SpliceAI) or acquiring or losing splicing site properties (Pangolin). The results for POLE and POLD1 were subsequently aggregated into an Excel file and uploaded as supplementary **Tables S5** and **S6**.

### GEMME Classification using a Gaussian Mixture Model

For classification with GEMME, we also adopted the procedure described in the ProteoCast framework [84]. For each protein, we modelled the distribution of GEMME scores using a three-component Gaussian Mixture Model (GMM). The model was fitted to the GEMME scores (the same score reported in the MAVISp “GEMME Score” column on the MAVISp csv file and in the original normPred_evolCombi output from GEMME), which represent the normalized evolutionary-energy estimates for all possible missense substitutions. The three GMM components were ordered by their mean scores, where more negative GEMME scores indicate stronger deleterious effects. Based on this order, we assigned a functional class to each component, following the ProteoCast nomenclature: i) i*mpactful*: component with the lowest mean (most deleterious score distribution), ii) *mild*: intermediate-mean component, and iii) *neutral*: highest-mean component. Each variant was then classified by maximum posterior probability, i.e., assigned to the class whose mixture component had the highest posterior for its GEMME score. The code is available at https://github.com/ELELAB/mavisp_accessory_tools.

### Downstream analyses and visualization of MAVISp data

We used the scripts and protocols from the downstream analysis toolkit of MAVISp (https://github.com/ELELAB/MAVISp_downstream_analysis) to facilitate the interpretation of the results and the selection of variants for further investigation. This included dot_plot and lolliplot visualization for MAVISp features and classes of mechanistic indicators, respectively. Additionally, barplots to compare the actual predicted values of changes in folding free energy by the different methods applied in the STABILITY module.

### Ranking of variants using MAVISp features

VUS and variants with conflicting evidence from ClinVar were prioritized using a feature-based ranking pipeline implemented in Python (https://github.com/ELELAB/MAVISp_downstream_analysis). The ranking was carried out after the application of the default discovery workflow of MAVISp based on AlphaMissense and DeMaSk. Each variant was scored across multiple structural and evolutionary features defined in a configuration file and extracted from the MAVISp output, including changes in binding free energies (i.e., local interactions), folding free energies (i.e., stability), changes upon phosphorylation, accuracy of the structural model (i.e., AlphaFold 2 pLDDT score), VEP scores and gnomAD genome/exome allele frequencies. Details on the features and weights are reported in the config files in the OSF repository. Quantitative features were min-max normalized, while categorical annotations were converted to ordinal scores. Each feature contributed to an overall composite score as a weighted sum of normalized feature values, where weights were specified in the configuration file. We gave higher weights to the mechanistic indicators, and among those, the highest weights were associated with local interactions because their changes in free energies generally have a narrower distribution with respect to the changes in folding free energy predicted by the STABILITY module. Variants were ranked in descending order of total score, and the top candidates were visualized with a feature heatmap to illustrate which features contributed most to each variant prioritization.

### Computational Re-classification of ClinVar VUS Using MAVISp-Derived Evidence

We evaluated whether MAVISp-derived mechanistic indicators could provide ACMG/AMP-compatible evidence for the interpretation of ClinVar variants of uncertain significance (VUS) in POLE and POLD1. The analysis followed the current ClinGen Variant Curation Expert Panel (VCEP) specification for polymerase proofreading genes, focusing on the computational evidence criteria PP3, BP4, and the structural/functional evidence adapted to a PM1-like category. We used the full MAVISp *ensemble-mode* annotations for POLE and POLD1 (this study) and retained all variants with available ClinVar interpretation. Variants annotated in ClinVar as uncertain significance, conflicting or not provided were classified as VUS and included for reevaluation. Pathogenic/likely pathogenic (P/LP) and benign/likely benign (B/LB) variants were used exclusively for threshold calibration and were not included in the final scoring. To generate quantitative thresholds for PP3/BP4, we extracted the VEP scores present in MAVISp (i.e., REVEL, AlphaMissense, EVE, GEMME and DeMaSk). Following ACMG/AMP guidelines, each predictor was calibrated using P/LP and B/LB variants. For each metric we computed: AUC, 25th percentile of pathogenic variants, 75th percentile of benign variants, Youden J optimal threshold, and Mann-Whitney U test to quantify effect-size separation. For each VUS, PP3 was assigned if a predictor exceeded (or, where relevant, fell below) the calibrated pathogenic threshold; BP4 was assigned if the value fell within the calibrated benign range. Following ClinGen guidance, any PP3 call overrides BP4, while the presence of any BP4 results in a BP4 assignment when PP3 is absent. Predictors that produced NaN values were ignored for that variant. We complemented PP3/BP4 with a class of MAVISp-derived PM1-like evidence, consistent with ClinGen specifications. A variant received PM1-like TRUE if any of the following MAVISp modules supported local or non-local damaging effect: FUNCTIONAL_SITES, LONG_RANGE, LOCAL_INTERACTIONS, STABILITY and PTM. For each VUS we reported: i) Individual PP3/BP4 status for five predictors, ii) Combined PP3/BP4 computational evidence, iii) PM1-like evidence flag, iv) Number of orthogonal computational lines of evidence. The scripts to reproduce this protocol are reported in https://github.com/ELELAB/MAVISp_downstream_analysis.

## Acknowledgements

Our research has been supported by Danmarks Grundforskningsfond (DNRF125), Novo Nordisk Fonden Bioscience and Basic Biomedicine (NNF20OC0065262) to the E.P. group. Part of the calculations have been supported by a EuroHPC Benchmark Access Grant (EHPC-BEN-2023B02-010) and a EuroHPC Regular Grant (EHPC-REG-2023R01-051) on Discoverer. This work was supported by Danish Data Science Academy, which is funded by the Novo Nordisk Foundation (NNF21SA0069429) and VILLUM FONDEN (40516); by The Assar Gabrielsson’s foundation and The Healthcare Board, Region ValJstra GolJtaland.

## Author Contributions

*Conceptualization*: EP *Data Curation*: MA, LB, KK, PSIB, AR, EP *Formal Analysis*: MA, EP *Investigation:* MA, AR, EP *Funding Acquisition:* EP, AR, MN, MA *Methodology*: MA, LB, KK, MT, EP *Project administration:* EP. *Resources*: EP, AR. *Software*: MA, LB, KK, MT, EP *Supervision*: EP. *Validation*: All the co-authors. *Visualization*: MA, EP. *Writing – Original Draft:* MA, EP. *Writing – Review and Editing*: All the coauthors.

## Declaration of generative AI and AI-assisted technologies in the writing process

During the preparation of this work the author(s) used OpenAI ChatGPT 5.0 to improve the language of the manuscript. After using this service, the authors reviewed and edited the content as needed and take full responsibility for the content of the published article.

## Data availability statement

All data and code, including input files and simulation trajectories, are freely available on OSF (https://osf.io/z8x4j) and the different GitHub repositories provided in the Materials and Methods Section. The aggregated data for POLE and POLD1 are also available as entries in the MAVISp database (https://services.healthtech.dtu.dk/services/MAVISp-1.0/).

## Supplementary material

**Supplementary Figure S1. Overview of the initial inputs for the MAVISp framework on POLE and POLD1**

A) Three-dimensional (3D) structure of POLE (1-2286) as retrieved from the AlphaFold (AF) Database (AFDB), and trimmed to 2-2286 residues for use in the simple mode calculation of the MAVISp framework, colored according to the pLDDT score, using the same color scale as the AFDB. Zn2+ ions have been modelled through superimposition with the experimental structure. B) 3D structure of POLD1 (1-1107) structure as retrieved by the AlphaFold (AF) Database (AFDB), and trimmed to 2-1107 residues for use in the simple mode calculation of the MAVISp framework, colored according to the pLDDT score, using the same color scale as AFDB. Zn2+ ion has been modelled through AlphaFill. C-D) Variant Annotations for POLE (C) and POLD1 (D) The UpSet plot showing the number of mutations in POLE analyzed with MAVISp grouped by source type. The plot displays the distribution of variants reported also in ClinVar, cBioPortal, or COSMIC against the total number of variants, here represented by the saturation mutagenesis scheme used in MAVISp.

**Supplementary Figure S2. Top 25 VUS from the ranking protocol applied to the MAVISp data** on variants with conflicting evidence from ClinVar for which MAVISp can identify structural alterations and VEPs predict damaging effects for POLE. All the variants have been ranked using the most important features provided by MAVISp with associated weights as explained in Materials and Methods. The corresponding dotplots are reported in Figure 6 for an overview of the effects and the individual predictions.

**Supplementary Figure S3. Top 25 VUS from the ranking protocol applied to the MAVISp data** on variants with conflicting evidence from ClinVar for which MAVISp can identify structural alterations and VEPs predict damaging effects for POLD1. All the variants have been ranked using the most important features provided by MAVISp with associated weights as explained in Materials and Methods. The corresponding dotplots are reported in Figure 6 for an overview of the effects and the individual predictions.

**Supplementary Figure S4. Contact analysis of exonuclease domain residues in POLE and POLD1.**

Contact analysis for residues I296 of POLE (A), F315 of POLD1 (B), G395 of POLD1 (C), L465 of POLE (D), I293 of POLE (E) along the POLE and POLD1 molecular dynamics trajectories, illustrating the occurrence and persistence of residue–residue interactions over time. Contact frequency is expressed as the number of simulation frames in which each interaction is observed.

**Supplementary Figure S5. Solvent Accessibility (SASA) Analysis of S314 in POLD1.**

(A) Time series of the relative solvent accessibility of the side-chain atoms of S314 calculated from MD simulation frames sampled every 10 ps during the evolution of the POLD1 system.

(B) Density distributions of SASA values for the side-chain atoms of POLD1 S314 computed across all the frames of the MD simulation.

**Supplementary Table S1.** Variants of POLE and POLD1 reported as pathogenic, likely pathogenic, benign, or likely benign in ClinVar at the date of 07/11/2025.

**Supplementary Table S2**. Calibrated thresholds for PP3/BP4 computational evidence in POLE and POLD1 exonuclease-domain variants.

This table reports the POLE/POLD1-specific calibration results for all variant effect predictors (VEPs) used to assign ACMG/AMP computational evidence (PP3 and BP4). Thresholds were derived using ClinVar pathogenic/likely pathogenic (P/LP) and benign/likely benign (B/LB) missense variants within the exonuclease domain (POLE: residues 268–471; POLD1: residues 304–533). For each predictor, we provide the 25th percentile of the pathogenic distribution (used as the PP3-supporting boundary), the 75th percentile of the benign distribution (BP4-supporting boundary), the ROC AUC, the optimal Youden J threshold, and the Mann–Whitney U test p-value. These thresholds were applied exclusively to ClinVar VUS in the exonuclease domain following ClinGen SVI recommendations for PP3/BP4 calibration.

**Supplementary Tables S3 and S4.** ACMG-based PP3, BP4, and PM1-like classifications for all POLE (S3) and POLD1 (S4) variants analyzed in MAVISp.

**Supplementary Table S5. Predictions with Pangolin of effects induced upon mutation on splicing sites**. We included the genomic coordinates of the mutation, which were provided as input to Pangolin, listed under the “genomic_coordinates” column. The table also provides the position within a 500-nucleotide window (default: ±500 bp) that may be affected by the variant, as specified in the “affected_nucleotide_position” column. The predicted types of splicing alterations caused by the variants are detailed in the “Δ_type” column. These include “splice gain”, where the nucleotide acquires splicing activity, and “splice loss”, where the nucleotide loses splicing activity. The “Δ_score” column reflects the difference between the reference (REF) and alternate (ALT) scores obtained from Pangolin for a given position. These scores represent the likelihood of a position acquiring or losing splicing site properties. The “REF_score” and *ALT_score* columns provide the computed probabilities that the position is a splice site based on the reference and alternate haplotypes. The table also includes the *gene_id* column, which lists the Ensembl gene code representing the gene containing the mutation, and the *transcript_id* column, which provides the Ensembl transcript code corresponding to the transcript affected by the mutation. Lastly, the *ref_seq_id* column includes the RefSeq identifier of the protein (if applicable) affected by the mutation, assuming the transcript encodes a protein.

**Supplementary Table S6 Predictions with SpliceAI**. We included the genomic coordinates of the mutation, which were provided as input to SpliceAI, listed under the “genomic_coordinates” column. The table also provides the position within a 500-nucleotide window (default: ±500 bp) that may be affected by the variant, as specified in the “affected_nucleotide_position” column. The predicted types of splicing alterations caused by the variants are detailed in the “Δ_type” column. These include “acceptor gain”, indicating that the nucleotide specified in the “affected_nucleotide_position” gains splicing acceptor activity; “donor gain”, where the nucleotide gains splicing donor activity; “acceptor loss”, where the nucleotide loses splicing acceptor activity; and “donor loss”, where the nucleotide loses splicing donor activity. The “Δ_score” column reflects the difference between the reference (REF) and alternate (ALT) scores obtained from SpliceAI for a given position. These scores represent the likelihood of a position becoming an acceptor or donor site or losing its acceptor or donor activity. The “REF_score” and *ALT_score* columns provide the computed probabilities of a given position being a splice acceptor or donor based on the reference and alternate haplotype sequences, respectively, as predicted by SpliceAI. The table also includes the *gene_id* column, which lists the Ensembl gene code representing the gene containing the mutation, and the *transcript_id* column, which provides the Ensembl transcript code corresponding to the transcript affected by the mutation. Lastly, the *ref_seq_id* column includes the RefSeq identifier of the protein (if applicable) affected by the mutation, assuming the transcript encodes a protein.

## References

1. Bębenek A, Ziuzia-Graczyk I. Fidelity of DNA replication—a matter of proofreading. Current Genetics. 2018. doi:10.1007/s00294-018-0820-1

2. Strauss JD, Pursell ZF. Replication DNA polymerases, genome instability and cancer therapies. NAR Cancer. 2023;5. doi:10.1093/narcan/zcad033

3. Mur P, García-Mulero S, del Valle J, Magraner-Pardo L, Vidal A, Pineda M, et al. Role of POLE and POLD1 in familial cancer. Genetics in Medicine. 2020;22. doi:10.1038/s41436-020-0922-2

4. Henninger EE, Pursell ZF. DNA polymerase ε and its roles in genome stability. IUBMB Life. 2014. doi:10.1002/iub.1276

5. Nicolas E, Golemis EA, Arora S. POLD1: Central mediator of DNA replication and repair, and implication in cancer and other pathologies. Gene. 2016. doi:10.1016/j.gene.2016.06.031

6. Prindle MJ, Loeb LA. DNA polymerase delta in dna replication and genome maintenance. Environmental and Molecular Mutagenesis. 2012. doi:10.1002/em.21745

7. Landrum MJ, Chitipiralla S, Brown GR, Chen C, Gu B, Hart J, et al. ClinVar: improvements to accessing data. Nucleic Acids Res. 2020;48: D835–D844. doi:10.1093/NAR/GKZ972

8. Arnaudi M, Utichi M, Degn K, Beltrame L, Scrima S, Krzesińska K, et al. MAVISp: A Modular Structure-Based Framework for Protein Variant Effects. BioRxiv. 2022. pp. 1–23. doi:10.1101/2022.10.22.513328

9. Viana-Errasti J, Marín R, García-Mulero S, Pons T, Terradas M, Capellá G, et al. Comparative Analysis of Somatic and Germline Polymerase Proofreading Deficiencies in Cancer: Molecular and Clinical Implications. Modern Pathology. 2025;38. doi:10.1016/j.modpat.2025.100843

10. Park VS, Pursell ZF. POLE proofreading defects: Contributions to mutagenesis and cancer. DNA Repair. 2019. doi:10.1016/j.dnarep.2019.02.007

11. Xing X, Kane DP, Bulock CR, Moore EA, Sharma S, Chabes A, et al. A recurrent cancer-associated substitution in DNA polymerase ε produces a hyperactive enzyme. Nat Commun. 2019;10. doi:10.1038/s41467-018-08145-2

12. Mur P, Viana-Errasti J, García-Mulero S, Magraner-Pardo L, Muñoz IG, Pons T, et al. Recommendations for the classification of germline variants in the exonuclease domain of POLE and POLD1. Genome Med. 2023;15. doi:10.1186/s13073-023-01234-y

13. Li H, Xie B, Zhou Y, Rahmeh A, Trusa S, Zhang S, et al. Functional roles of p12, the fourth subunit of human DNA polymerase δ. Journal of Biological Chemistry. 2006;281. doi:10.1074/jbc.M600322200

14. Meng X, Zhou Y, Lee EYC, Lee MYWT, Frick DN. The p12 subunit of human polymerase Δ modulates the rate and fidelity of DNA synthesis. Biochemistry. 2010;49. doi:10.1021/bi100042b

15. Lin SHS, Wang X, Zhang S, Zhang Z, Lee EYC, Lee MYWT. Dynamics of enzymatic interactions during short flap human Okazaki fragment processing by two forms of human DNA polymerase δ. DNA Repair (Amst). 2013;12. doi:10.1016/j.dnarep.2013.08.008

16. Cheng J, Novati G, Pan J, Bycroft C, Žemgulytė A, Applebaum T, et al. Accurate proteome-wide missense variant effect prediction with AlphaMissense. Science. 2023;381: eadg7492. doi:10.1126/science.adg7492

17. Frazer J, Notin P, Dias M, Gomez A, Min JK, Brock K, et al. Disease variant prediction with deep generative models of evolutionary data. Nature. 2021;599: 91–95. doi:10.1038/s41586-021-04043-8

18. Ioannidis NM, Rothstein JH, Pejaver V, Middha S, McDonnell SK, Baheti S, et al. REVEL: An Ensemble Method for Predicting the Pathogenicity of Rare Missense Variants. Am J Hum Genet. 2016;99: 877–885. doi:10.1016/j.ajhg.2016.08.016

19. Laine E, Karami Y, Carbone A. GEMME: A Simple and Fast Global Epistatic Model Predicting Mutational Effects. Mol Biol Evol. 2019;36: 2604–2619. doi:10.1093/molbev/msz179

20. Zhou JC, Janska A, Goswami P, Renault L, Ali FA, Kotecha A, et al. CMG-Pol epsilon dynamics suggests a mechanism for the establishment of leading-strand synthesis in the eukaryotic replisome. Proc Natl Acad Sci U S A. 2017;114. doi:10.1073/pnas.1700530114

21. Goswami P, Abid Ali F, Douglas ME, Locke J, Purkiss A, Janska A, et al. Structure of DNA-CMG-Pol epsilon elucidates the roles of the non-catalytic polymerase modules in the eukaryotic replisome. Nat Commun. 2018;9. doi:10.1038/s41467-018-07417-1

22. Jones ML, Baris Y, Taylor MRG, Yeeles JTP. Structure of a human replisome shows the organisation and interactions of a DNA replication machine. EMBO J. 2021;40. doi:10.15252/embj.2021108819

23. Tiberti M, Terkelsen T, Degn K, Beltrame L, Canter Cremers T, da Piedade I, et al. MutateX: an automated pipeline for in silico saturation mutagenesis of protein structures and structural ensembles. Brief Bioinform. 2022; 1–16. doi:10.1093/bib/bbac074

24. Campbell BB, Light N, Fabrizio D, Zatzman M, Fuligni F, de Borja R, et al. Comprehensive Analysis of Hypermutation in Human Cancer. Cell. 2017;171. doi:10.1016/j.cell.2017.09.048

25. Zou Y, Liu FY, Liu H, Wang F, Li W, Huang MZ, et al. Frequent POLE1 p.S297F mutation in Chinese patients with ovarian endometrioid carcinoma. Mutation Research - Fundamental and Molecular Mechanisms of Mutagenesis. 2014;761. doi:10.1016/j.mrfmmm.2014.01.003

26. Herzog M, Alonso-Perez E, Salguero I, Warringer J, Adams DJ, Jackson SP, et al. Mutagenic mechanisms of cancer-associated DNA polymerase alleles. Nucleic Acids Res. 2021;49. doi:10.1093/nar/gkab160

27. Barbari SR, Kane DP, Moore EA, Shcherbakova P V. Functional analysis of cancer-associated DNA polymerase ε variants in Saccharomyces cerevisiae. G3: Genes, Genomes, Genetics. 2018;8. doi:10.1534/g3.118.200042

28. Labrousse G, Perre P Vande, Parra G, Jaffrelot M, Leroy L, Chibon F, et al. The hereditary N363K POLE exonuclease mutant extends PPAP tumor spectrum to glioblastomas by causing DNA damage and aneuploidy in addition to increased mismatch mutagenicity. NAR Cancer. 2023;5. doi:10.1093/narcan/zcad011

29. Dahl JM, Thomas N, Tracy MA, Hearn BL, Perera L, Kennedy SR, et al. Probing the mechanisms of two exonuclease domain mutators of DNA polymerase ε. Nucleic Acids Res. 2022;50. doi:10.1093/nar/gkab1255

30. Roske JJ, Yeeles JTP. Structural basis for processive daughter-strand synthesis and proofreading by the human leading-strand DNA polymerase Pol ε. Nat Struct Mol Biol. 2024. doi:10.1038/s41594-024-01370-y

31. Ostroverkhova D, Tyryshkin K, Beach AK, Moore EA, Masoudi-Sobhanzadeh Y, Barbari SR, et al. DNA polymerase ε and δ variants drive mutagenesis in polypurine tracts in human tumors. Cell Rep. 2024;43. doi:10.1016/j.celrep.2023.113655

32. Rohlin A, Zagoras T, Nilsson S, Lundstam U, Wahlström J, Hultén L, et al. A mutation in POLE predisposing to a multi-tumour phenotype. Int J Oncol. 2014;45: 77–81. doi:10.3892/ijo.2014.2410

33. Djursby M, Madsen MB, Frederiksen JH, Berchtold LA, Therkildsen C, Willemoe GL, et al. New Pathogenic Germline Variants in Very Early Onset and Familial Colorectal Cancer Patients. Front Genet. 2020;11. doi:10.3389/fgene.2020.566266

34. Hamzaoui N, Alarcon F, Leulliot N, Guimbaud R, Buecher B, Colas C, et al. Genetic, structural, and functional characterization of POLE polymerase proofreading variants allows cancer risk prediction. Genetics in Medicine. 2020;22. doi:10.1038/s41436-020-0828-z

35. Herr AJ, Ogawa M, Lawrence NA, Williams LN, Eggington JM, Singh M, et al. Mutator suppression and escape from replication error-induced extinction in yeast. PLoS Genet. 2011;7. doi:10.1371/journal.pgen.1002282

36. Brown JA, Suo Z. Unlocking the sugar “steric gate” of DNA polymerases. Biochemistry. 2011;50. doi:10.1021/bi101915z

37. Jubb HC, Higueruelo AP, Ochoa-Montaño B, Pitt WR, Ascher DB, Blundell TL. Arpeggio: A Web Server for Calculating and Visualising Interatomic Interactions in Protein Structures. J Mol Biol. 2017;429: 365–371. doi:10.1016/j.jmb.2016.12.004

38. Murphy K, Darmawan H, Schultz A, Da Silva EF, Reha-Krantz LJ. A method to select for mutator DNA polymerase δs in Saccharomyces cerevisiae. Genome. 2006;49. doi:10.1139/G05-106

39. COSMIC, Catalogue Of Somatic Mutations In Cancer, https://cancer.sanger.ac.uk.

40. Gao J, Aksoy BA, Dogrusoz U, Dresdner G, Gross B, Sumer SO, et al. Integrative analysis of complex cancer genomics and clinical profiles using the cBioPortal. Sci Signal. 2013;6: pl1. doi:10.1126/scisignal.2004088

41. Hornbeck P V., Zhang B, Murray B, Kornhauser JM, Latham V, Skrzypek E. PhosphoSitePlus, 2014: mutations, PTMs and recalibrations. Nucleic Acids Res. 2015;43: D512–D520. doi:10.1093/nar/gku1267

42. Bah A, Forman-Kay JD. Modulation of intrinsically disordered protein function by post-translational modifications. Journal of Biological Chemistry. 2016;291: 6696–705. doi:10.1074/jbc.R115.695056

43. Tiberti M, Di Leo L, Vistesen MV, Kuhre RS, Cecconi F, De Zio D, et al. The Cancermuts software package for the prioritization of missense cancer variants: a case study of AMBRA1 in melanoma. Cell Death Dis. 2022;13: 872. doi:10.1038/s41419-022-05318-2

44. Hopkins JJ, Wakeling MN, Johnson MB, Flanagan SE, Laver TW. REVEL Is Better at Predicting Pathogenicity of Loss-of-Function than Gain-of-Function Variants. Chen J-M, editor. Hum Mutat. 2023;2023: 1–6. doi:10.1155/2023/8857940

45. Jumper J, Evans R, Pritzel A, Green T, Figurnov M, Ronneberger O, et al. Highly accurate protein structure prediction with AlphaFold. Nature. 2021;596: 583–589. doi:10.1038/S41586-021-03819-2

46. Berman HM, Westbrook J, Feng Z, Gilliland G, Bhat TN, Weissig H, et al. The Protein Data Bank. Nucleic Acids Res. 2000;28: 235–242. doi:10.1093/NAR/28.1.235

47. Baranovskiy AG, Gu J, Babayeva ND, Kurinov I, Pavlov YI, Tahirov TH. Crystal structure of the human Polε B-subunit in complex with the C-terminal domain of the catalytic subunit. Journal of Biological Chemistry. 2017;292: 15717–15730. doi:10.1074/jbc.M117.792705

48. Lancey C, Tehseen M, Raducanu VS, Rashid F, Merino N, Ragan TJ, et al. Structure of the processive human Pol δ holoenzyme. Nat Commun. 2020;11. doi:10.1038/s41467-020-14898-6

49. Jozwiakowski SK, Kummer S, Gari K. Human DNA polymerase delta requires an iron–sulfur cluster for high-fidelity DNA synthesis. Life Sci Alliance. 2019;2. doi:10.26508/lsa.201900321

50. Calderone A, Castagnoli L, Cesareni G. Mentha: A resource for browsing integrated protein-interaction networks. Nat Methods. 2013;10: 690–691. doi:10.1038/nmeth.2561

51. Jenkyn-Bedford M, Jones ML, Baris Y, Labib KPM, Cannone G, Yeeles JTP, et al. A conserved mechanism for regulating replisome disassembly in eukaryotes. Nature. 2021;600: 743–747. doi:10.1038/s41586-021-04145-3

52. He Q, Wang F, Yao NY, O’Donnell ME, Li H. Structures of the human leading strand Polε–PCNA holoenzyme. Nature Communications. 2024;15. doi:10.1038/s41467-024-52257-x

53. Zhu W, Shenoy A, Kundrotas P, Elofsson A. Evaluation of AlphaFold-Multimer prediction on multi-chain protein complexes. Cowen L, editor. Bioinformatics. 2023;39. doi:10.1093/bioinformatics/btad424

54. Burke DF, Bryant P, Barrio-Hernandez I, Memon D, Pozzati G, Shenoy A, et al. Towards a structurally resolved human protein interaction network. Nat Struct Mol Biol. 2023;30. doi:10.1038/s41594-022-00910-8

55. Delgado J, Radusky LG, Cianferoni D, Serrano L. FoldX 5.0: Working with RNA, small molecules and a new graphical interface. Bioinformatics. 2019; 1–2. doi:10.1093/bioinformatics/btz184

56. Pasi M, Tiberti M, Arrigoni A, Papaleo E. xPyder: a PyMOL plugin to analyze coupled residues and their networks in protein structures. J Chem Inf Model. 2012;279: 1–6. doi:10.1021/ci300213c

57. Abraham MJ, Murtola T, Schulz R, Páll S, Smith JC, Hess B, et al. GROMACS: High performance molecular simulations through multi-level parallelism from laptops to supercomputers. SoftwareX. 2015;2: 19–25. doi:10.1016/j.softx.2015.06.001

58. Huang J, Rauscher S, Nawrocki G, Ran T, Feig M, de Groot BL, et al. CHARMM36m: an improved force field for folded and intrinsically disordered proteins. Nat Methods. 2017;14: 71–73. doi:10.1038/nmeth.4067

59. Hess B, Bekker H, Berendsen HJC, Fraaije JGEM. LINCS: A linear constraint solver for molecular simulations. J Comput Chem. 1997;18: 1463–1472. doi:10.1002/(SICI)1096-987X(199709)18:12<1463::AID-JCC4>3.0.CO;2-H

60. Essmann U, Perera L, Berkowitz ML, Darden T, Lee H, Pedersen LG. A smooth particle mesh Ewald method. J Chem Phys. 1995;103: 8577–8593. doi:10.1063/1.470117

61. Darden T, York D, Pedersen L. Particle mesh Ewald: An *N* log(*N*) method for Ewald sums in large systems. J Chem Phys. 1993;98: 10089–10092. doi:10.1063/1.464397

62. Berendsen HJC, Postma JPM, van Gunsteren WF, DiNola A, Haak JR. Molecular dynamics with coupling to an external bath. J Chem Phys. 1984;81: 3684–3690. doi:10.1063/1.448118

63. Nosé S, Klein ML. Constant pressure molecular dynamics for molecular systems. Mol Phys. 1983;50: 1055–1076. doi:10.1080/00268978300102851

64. Blaabjerg LM, Kassem MM, Good LL, Jonsson N, Cagiada M, Johansson KE, et al. Rapid protein stability prediction using deep learning representations. Elife. 2023;12: e82593. doi:10.7554/eLife.82593

65. Sora V, Tiberti M, Robbani SM, Rubin J, Papaleo E. PyInteraph2 and PyInKnife2 to analyze networks in protein structural ensembles. bioRxiv. 2020. doi:10.1101/2020.11.22.381616

66. Barlow KA, Ó Conchúir S, Thompson S, Suresh P, Lucas JE, Heinonen M, et al. Flex ddG: Rosetta Ensemble-Based Estimation of Changes in Protein-Protein Binding Affinity upon Mutation. Journal of Physical Chemistry B. 2018;122: 5389–5399. doi:10.1021/acs.jpcb.7b11367

67. Alford RF, Leaver-Fay A, Jeliazkov JR, O’Meara MJ, DiMaio FP, Park H, et al. The Rosetta All-Atom Energy Function for Macromolecular Modeling and Design. J Chem Theory Comput. 2017;13: 3031–3048. doi:10.1021/acs.jctc.7b00125

68. Raimondi D, Orlando G, Pancsa R, Khan T, Vranken WF. Exploring the Sequence-based Prediction of Folding Initiation Sites in Proteins. Sci Rep. 2017;7: 8826. doi:10.1038/s41598-017-08366-3

69. Sora V, Otamendi Laspiur A, Degn K, Arnaudi M, Utichi M, Beltrame L, et al. RosettaDDGPrediction for high-throughput mutational scans: from stability to binding. Protein Science. 2023;32: e4527. doi:10.1002/pro.4527

70. Shapovalov M V., Dunbrack RL. A smoothed backbone-dependent rotamer library for proteins derived from adaptive kernel density estimates and regressions. Structure. 2011;19: 844–858. doi:10.1016/j.str.2011.03.019

71. Song Y, Tyka M, Leaver-Fay A, Thompson J, Baker D. Structure-guided forcefield optimization. Proteins: Structure, Function and Bioinformatics. 2011;79: 1898–1909. doi:10.1002/prot.23013

72. O’Meara MJ, Leaver-Fay A, Tyka MD, Stein A, Houlihan K, Dimaio F, et al. Combined covalent-electrostatic model of hydrogen bonding improves structure prediction with Rosetta. J Chem Theory Comput. 2015;11: 609–622. doi:10.1021/ct500864r

73. Hubbard SJ, Thornton JM,. NACCESS. 1993. Available: Department of Biochemistry and Molecular Biology, University College London

74. Kumar M, Gouw M, Michael S, Sámano-Sánchez H, Pancsa R, Glavina J, et al. ELM-the eukaryotic linear motif resource in 2020. Nucleic Acids Res. 2020. doi:10.1093/nar/gkz1030

75. Tan ZW, Guarnera E, Tee WV, Berezovsky IN. AlloSigMA 2: Paving the way to designing allosteric effectors and to exploring allosteric effects of mutations. Nucleic Acids Res. 2020;48: W116–W124. doi:10.1093/NAR/GKAA338

76. Le Guilloux V, Schmidtke P, Tuffery P. Fpocket: An open source platform for ligand pocket detection. BMC Bioinformatics. 2009;10: 1–11. doi:10.1186/1471-2105-10-168/TABLES/1

77. Bhattacharyya M, Ghosh S, Vishveshwara S. Protein Structure and Function: Looking through the Network of Side-Chain Interactions. Curr Protein Pept Sci. 2016;17: 4–25. doi:10.2174/1389203716666150923105727

78. Tiberti M, Invernizzi G, Lambrughi M, Inbar Y, Schreiber G, Papaleo E. PyInteraph: A framework for the analysis of interaction networks in structural ensembles of proteins. J Chem Inf Model. 2014;54: 1537–1551. doi:10.1021/ci400639r

79. Krzesińska K, Degn K, Llorente A, Giannakopoulou E, Tiberti M, Papaleo E. Deciphering Long-Range Effects of Mutations: An Integrated Approach Using Elastic Network Models and Protein Structure Networks. J Mol Biol. 2025;437: 169359. doi:10.1016/J.JMB.2025.169359

80. Dijkstra EW. A note on two problems in connexion with graphs. Numer Math (Heidelb). 1959;1: 269–271. doi:10.1007/BF01386390

81. Hagberg AA, Schult DA, Swart PJ. Exploring network structure, dynamics, and function using NetworkX. 7th Python in Science Conference (SciPy 2008). 2008; 11–15. doi:http://conference.scipy.org/proceedings/SciPy2008/paper_2

82. Jaganathan K, Kyriazopoulou Panagiotopoulou S, McRae JF, Darbandi SF, Knowles D, Li YI, et al. Predicting Splicing from Primary Sequence with Deep Learning. Cell. 2019;176. doi:10.1016/j.cell.2018.12.015

83. Zeng T, Li YI. Predicting RNA splicing from DNA sequence using Pangolin. Genome Biol. 2022;23. doi:10.1186/s13059-022-02664-4

84. Abakarova M, Freiberger MI, Liehrmann A, Rera M, Laine E. Proteome-wide Prediction of the Functional Impact of Missense Variants with ProteoCast. 2025. doi:10.1101/2025.02.09.637326

