## Supplementary Figure 1 for "Interpreting the Effects of DNA Polymerase Variants at the Structural Level Using MAVISp and Molecular Dynamics Simulations"

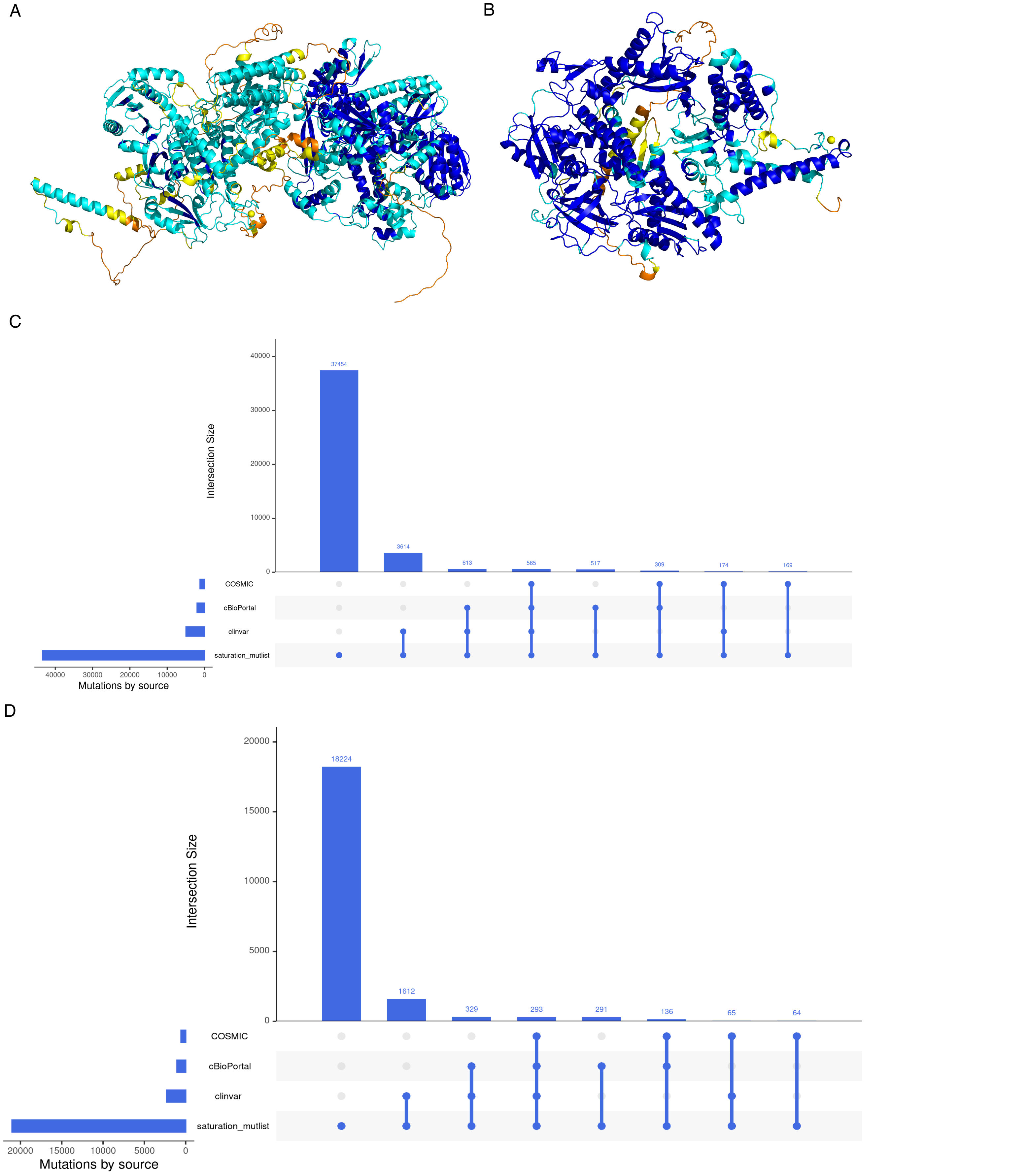


**FigureS1 Overview of the initial inputs for the MAVISp framework on POLE and POLD1** A) Three-dimensional (3D) structure of POLE (1-2286) as retrieved from the AlphaFold (AF) Database (AFDB), and trimmed to 2-2286 residues for use in the simple mode calculation of the MAVISp framework, colored according to the pLDDT score, using the same color scale as the AFDB. Zn2+ ions have been modelled through superimposition with the experimental structure. B) 3D structure of POLD1 (1-1107) structure as retrieved by the AlphaFold (AF) Database (AFDB), and trimmed to 2-1107 residues for use in the simple mode calculation of the MAVISp framework, colored according to the pLDDT score, using the same color scale as AFDB. Zn2+ ion has been modelled through AlphaFill. C-D) Variant Annotations for POLE (C) and POLD1 (D) The UpSet plot showing the number of mutations in POLE analyzed with MAVISp grouped by source type. The plot displays the distribution of variants reported also in ClinVar, cBioPortal, or COSMIC against the total number of variants, here represented by the saturation mutagenesis scheme used in MAVISp.
