## Supplementary Figure 2 for "Interpreting the Effects of DNA Polymerase Variants at the Structural Level Using MAVISp and Molecular Dynamics Simulations"

**Figure S2. Top 25 VUS from the ranking protocol applied to the MAVISp data** on variants with conflicting evidence from ClinVar for which MAVISp can identify structural alterations and VEPs predict damaging effects for POLE. All the variants have been ranked using the most important features provided by MAVISp with associated weights as explained in Materials and Methods**.** The corresponding dotplots are reported in Figure 6 for an overview of the effects and the individual predictions.

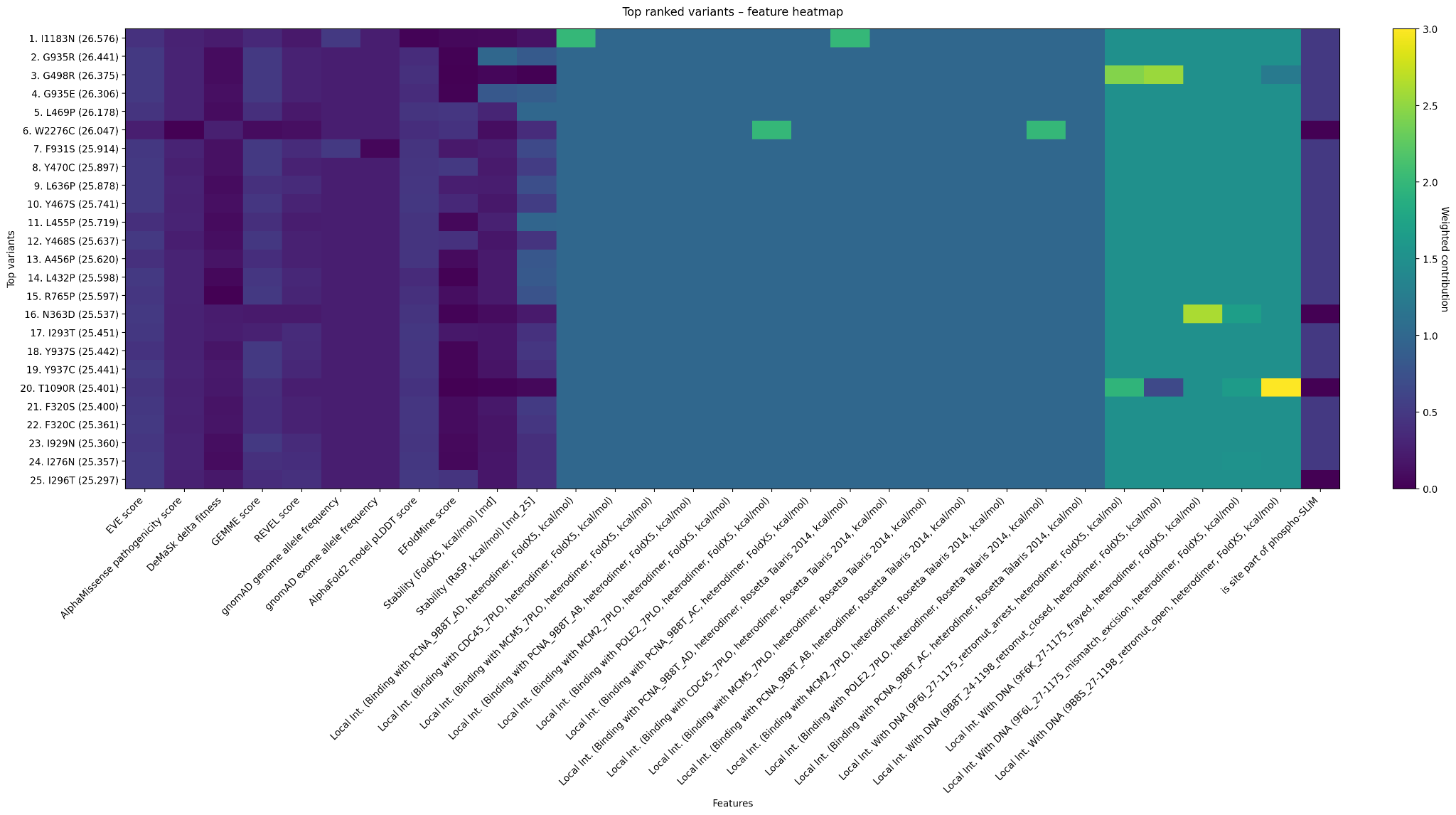
