## Supplementary Figure 4 for "Interpreting the Effects of DNA Polymerase Variants at the Structural Level Using MAVISp and Molecular Dynamics Simulations"

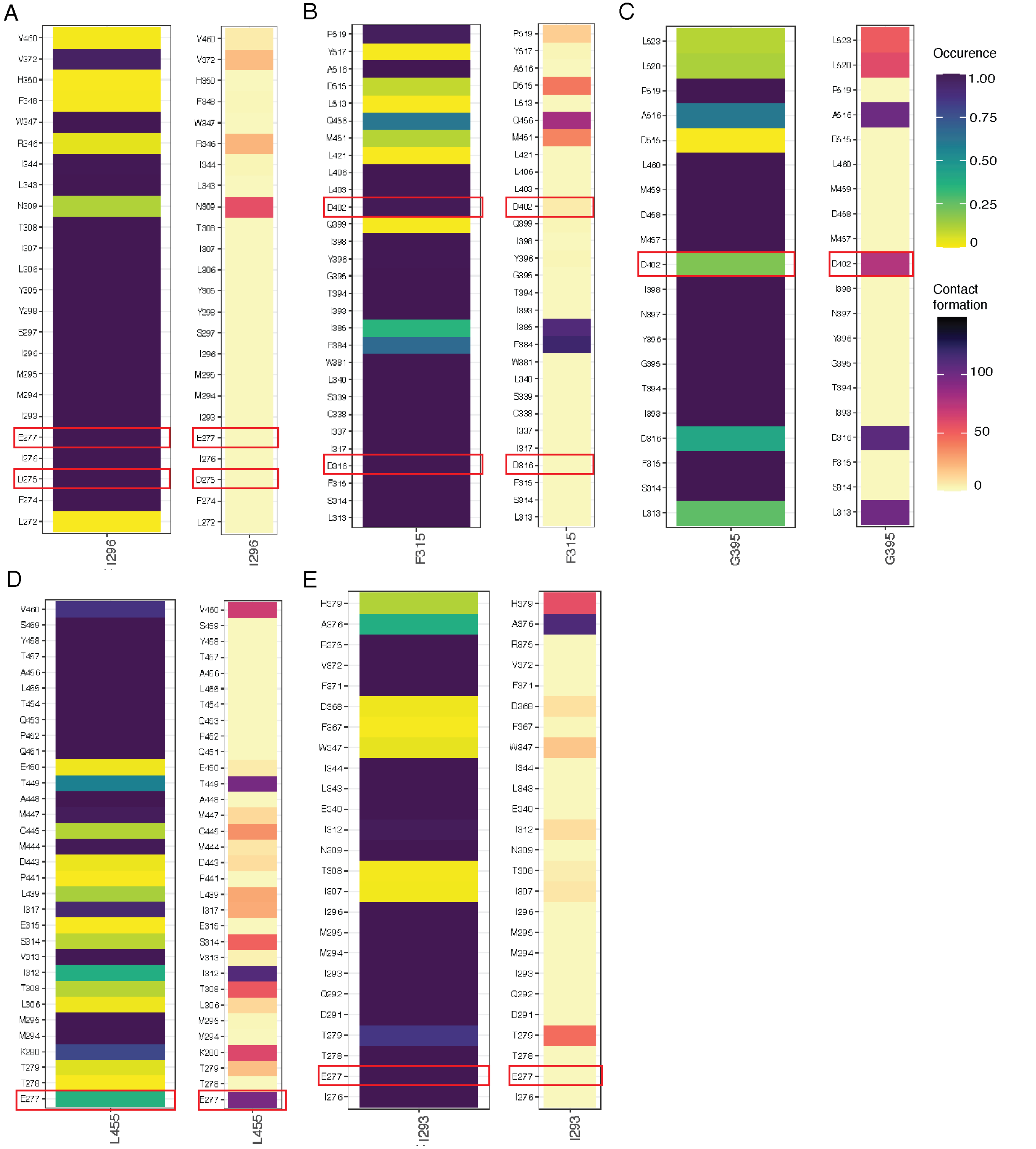


**Supplementary Figure S4. Contact analysis of exonuclease domain residues in POLE and POLD1.** Contact analysis for residues I296 of POLE (A), F315 of POLD1 (B), G395 of POLD1 (C), L465 of POLE (D) and I293 of POLE (E) along the POLE and POLD1 molecular dynamics trajectories, illustrating the occurrence and persistence of residue–residue interactions over time. Contact frequency is expressed as the number of simulation frames in which each interaction is observed. Relevant residues establishing contacts are highlighted in red.
