## Supplementary Figure 5 for "Interpreting the Effects of DNA Polymerase Variants at the Structural Level Using MAVISp and Molecular Dynamics Simulations"

A

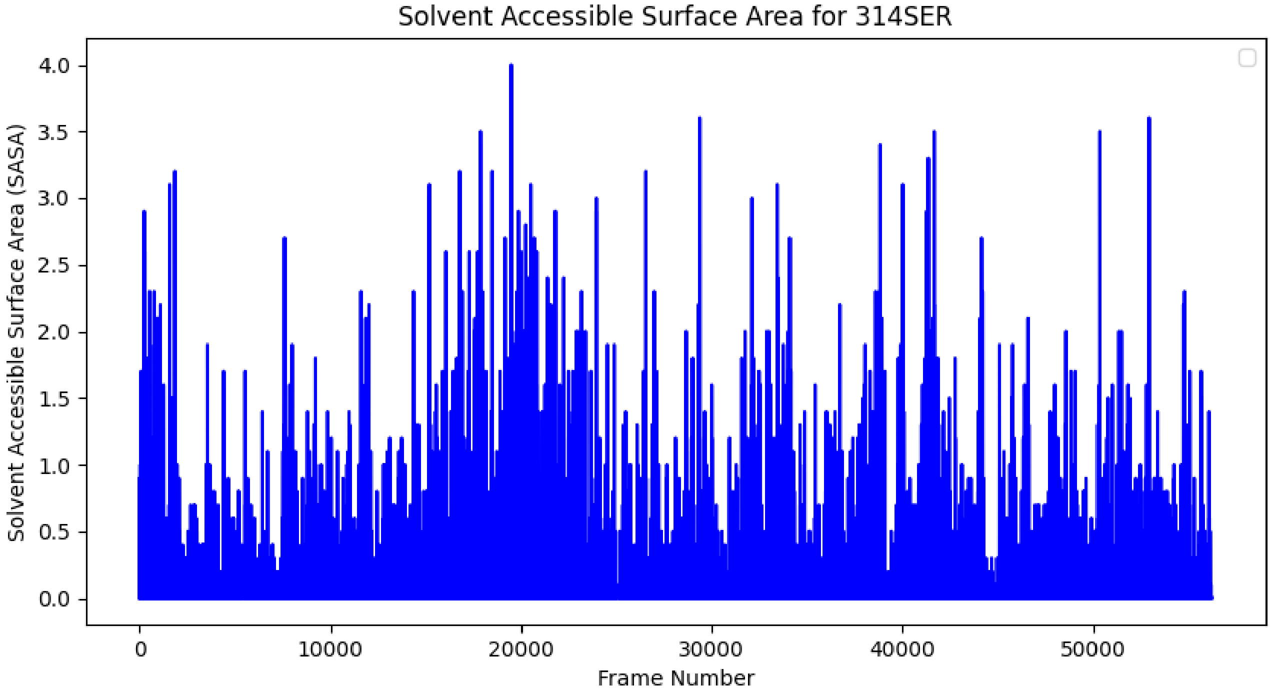


B

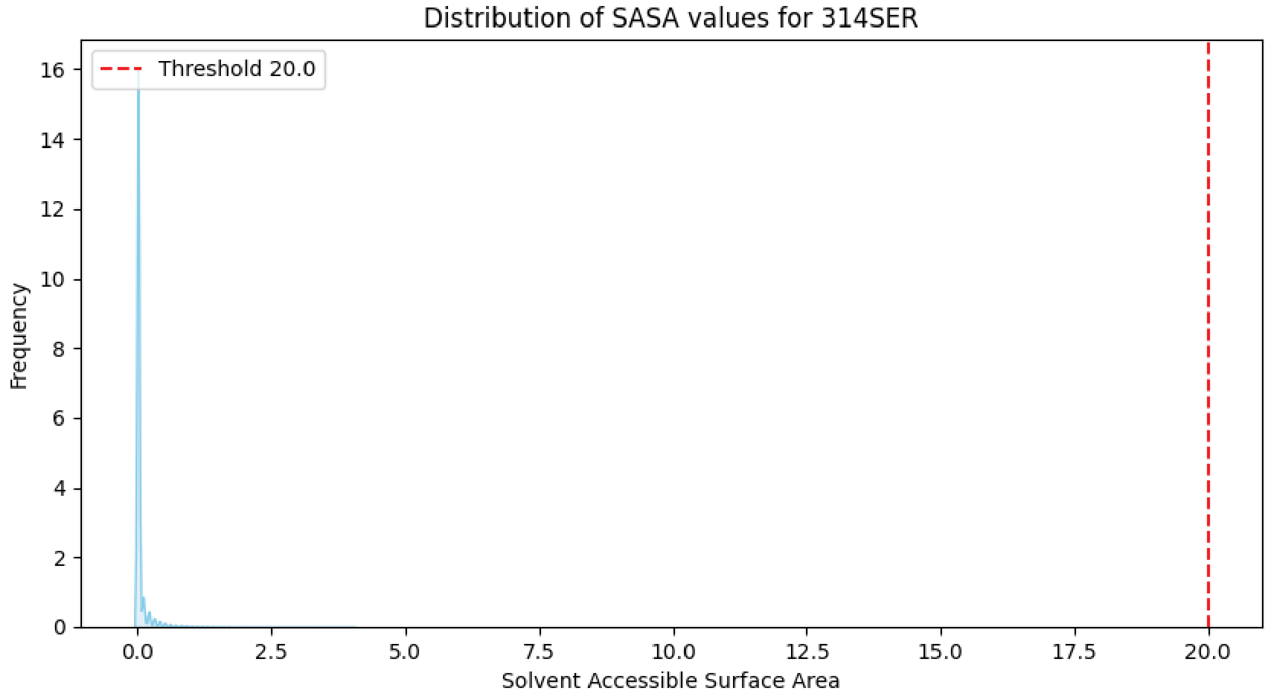


**Supplementary Figure S5. Solvent Accessibility (SASA) Analysis of S314 in POLD1.** (A) Time series of the relative solvent accessibility of the side-chain atoms of S314 calculated from MD simulation frames sampled every 10 ps during the evolution of the POLD1 system. (B) Density distributions of SASA values for the side-chain atoms of S314 computed across all the frames of the MD simulation.
